# Inhibition of cholinergic interneurons potentiates corticostriatal transmission in D1 receptor-expressing medium-sized spiny neurons and restores motor learning in parkinsonian condition

**DOI:** 10.1101/2021.07.07.451477

**Authors:** Gwenaelle Laverne, Jonathan Pesce, Ana Reynders, Christophe Melon, Lydia Kerkerian-Le Goff, Nicolas Maurice, Corinne Beurrier

## Abstract

Striatal cholinergic interneurons (CINs) respond to salient or reward prediction-related stimuli after conditioning with brief pauses in their activity, implicating them in learning and action selection. This pause is lost in animal models of Parkinson’s disease. How this signal regulates the functioning of the striatum remains an open question. To address this issue, we examined the impact of CIN firing inhibition on glutamatergic transmission between the cortex and the medium-sized spiny projection neurons expressing dopamine D1 receptors (D1 MSNs). Brief interruption of CIN activity had no effect in control condition whereas it increased glutamatergic responses in D1 MSNs after nigrostriatal dopamine denervation. This potentiation was dependent upon M4 muscarinic receptor and protein kinase A. Decreasing CIN firing by opto/chemogenetic strategies *in vivo* rescued long-term potentiation in some MSNs and alleviated motor learning deficits in parkinsonian mice. Taken together, our findings demonstrate that the control exerted by CINs on corticostriatal transmission and striatal-dependent motor-skill learning depends on the integrity of dopaminergic inputs.

## INTRODUCTION

Voluntary movement and action planning depend on the balanced activity of two distinct populations of GABAergic projection neurons in the striatum called medium-sized spiny neurons (MSNs). MSNs expressing D1 dopamine (DA) receptor (D1 MSNs) project to the basal ganglia output nuclei, forming the so-called direct pathway, whereas MSNs expressing D2 receptors (D2 MSNs) primarily project to the globus pallidus, forming the first link of the indirect pathway (Albin et al., 1989; DeLong, 1990). MSN firing relies on excitatory inputs from the cerebral cortex and the thalamus and is modulated by DA afferents from the substantia nigra pars compacta. DA is expected to induce motor activation by simultaneously activating D1 MSNs to promote selected actions and depressing D2 MSNs to suppress competing actions. In Parkinson’s disease (PD), the loss of nigrostriatal DA neurons results in an imbalance in favor of the indirect pathway, the activities of D1 MSNs and D2 MSNs being inappropriately decreased and increased, respectively (DeLong, 1990; Mallet et al., 2005; Smith et al., 1998). These cell-specific changes are thought to underlie most parkinsonian deficits, including slowness of movement.

In addition to DA, the striatum exhibits a rich cholinergic innervation and expresses high levels of acetylcholine (ACh), muscarinic receptors (mAChRs) and other markers related to ACh. mAChRs are distributed on the two populations of MSNs, on interneurons and on synaptic terminals, including glutamatergic axons (Goldberg et al., 2012). At the post-synaptic level, the Gαq-coupled M1 mAChR is present on both D1 and D2 MSNs while the Gαi-coupled M4 mAChR is preferentially expressed by D1 MSNs (Hersch et al., 1994; Yan et al., 2001). The effects of mAChR activation have been mostly investigated using application of exogenous agonists in conjunction with whole-cell patch-clamp recording *ex vivo*. This approach has revealed a variety of effects on MSN intrinsic excitability and excitatory synaptic transmission depending on the type of mAChR targeted (Abudukeyoumu et al., 2019; Zhai et al., 2018). However, sustained application of specific receptor agonists does not recapitulate the spatiotemporal dynamics of endogenous ACh.

In the striatum, cholinergic interneurons (CINs) are the main source of ACh. Despite their small number (1-2% of striatal cells), CINs harbor dense terminal fields that contact both populations of MSNs and are therefore well positioned to regulate striatal outflow. CINs transiently respond to motivationally relevant events with a brief pause in their tonic firing and are, for this reason, considered as key players in learning and decision-making (Aosaki et al., 1994a; Apicella et al.; Goldberg and Reynolds, 2011). The pause is mostly synchronized in the CIN population such as it might efficiently translate into global reduction of striatal ACh level (Aosaki et al., 1995). In PD, CINs lose their ability to pause and inhibition of their discharge using opto- or chemogenetic tools improves motor performances in mouse models of PD (Aosaki et al., 1994b; Maurice et al., 2015; Tanimura et al., 2019; Ztaou et al., 2016). A normalization of D2 MSN activity via their thalamic inputs might partly underlie this beneficial effect (Tanimura et al., 2019). On the other hand, we have shown that CIN opto-inhibition in parkinsonian mice had a selective impact on the component of the complex cortically-evoked neuronal responses in the substantia nigra pars reticulata mediated by the direct pathway (Maurice et al., 2015). This suggests that CINs also act on the trans-striatal processing of cortical information through D1 MSNs. However, how a synchronous inhibition of CIN activity shapes the dynamics of corticostriatal processing in these neurons in control and PD condition remains largely unexplored. To fill this gap, we used optogenetic inhibition of CIN firing in conjunction with recordings of synaptic responses triggered by cortical stimulation in genetically identified D1 MSNs in mouse striatal slices. After showing that inhibition of CIN firing triggered a potentiation of corticostriatal transmission in D1 MSNs in parkinsonian mice, but not in non-lesioned mice, we further explored the mechanisms underlying this effect. We also investigated whether optogenetic and chemogenetic inhibition of CINs *in vivo* could interfere with alterations in long-term potentiation at corticostriatal synapses and motor learning in parkinsonian condition. Our results demonstrate that, in parkinsonian state, the control exerted by CINs on corticostriatal transmission is altered in D1 MSNs via M4 mAChR-mediated mechanisms. Abnormal signaling by CINs also contributes to dysfunctional long-term plasticity at corticostriatal synapses and to disease symptoms.

## RESULTS

All mice used in this study were 2 to 4 months old at the time of experiments. Summary statistics (median and interquartile range, means and SEM), associated numbers of cells and mice, and p values for each statistical comparison are shown for all datasets in supplementary tables.

### Opto-inhibition of CINs induces a potentiation of corticostriatal synaptic transmission in D1 MSNs from 6-OHDA mice

In *ex vivo* brain slices, application of brief light stimulation (150 ms at 585 nm) induced a complete and reliable inhibition of the firing of halorhodopsin(eNpHR)-EYFP-expressing CINs (Figure S1). To determine whether such inhibition modulates corticostriatal synaptic transmission in control and PD-like condition, we recorded excitatory post-synaptic potentials (EPSPs) evoked by electrical stimulation of cortical fibers in D1 MSNs identified by tdTomato expression in triple-transgenic ChAT^cre/wt^; Rosa^NpHr/wt^; D1-tdTomato^+/−^ mice (see Methods for details) (Figure 1A). We found that the amplitude of the EPSPs, as well as the paired-pulse ratio (PPR, 50 ms interstimulus interval), were not significantly altered by CIN opto-inhibition in non-lesioned mice (Figure 1B and Table S1). These parameters were also not modified when CIN opto-inhibition was performed in the presence of neostigmine (Figure S2A and Table S2), an inhibitor of acetylcholinesterase known to increase extracellular ACh, which per se decreased EPSP amplitude and increased PPR (Figure S2B). Therefore, the lack of modulation of EPSP amplitude by CIN opto-inhibition was not due to insufficient cholinergic tone in slices. Consistently, opto-inhibition of CINs performed in anesthetized mice in which ACh tone is not altered by the slicing procedure did not impact the cortically-evoked EPSPs (Figure S3 and Table S2). In sharp contrast, CIN opto-inhibition significantly increased EPSP amplitude in D1 MSNs from 6-OHDA mice without affecting PPR (Figure 1C and Table S1). This potentiation was not observed in D1 MSNs recorded from littermate 6-OHDA mice that do not express Cre recombinase in cholinergic neurons, excluding non-specific light-induced effects (Figure S4). Together, these findings show that a brief interruption of CIN firing potentiates corticostriatal synaptic transmission in D1 MSNs after striatal DA denervation, suggesting that abnormal cholinergic signaling affects corticostriatal information processing through the direct pathway in PD-like condition.

**Figure 1:**
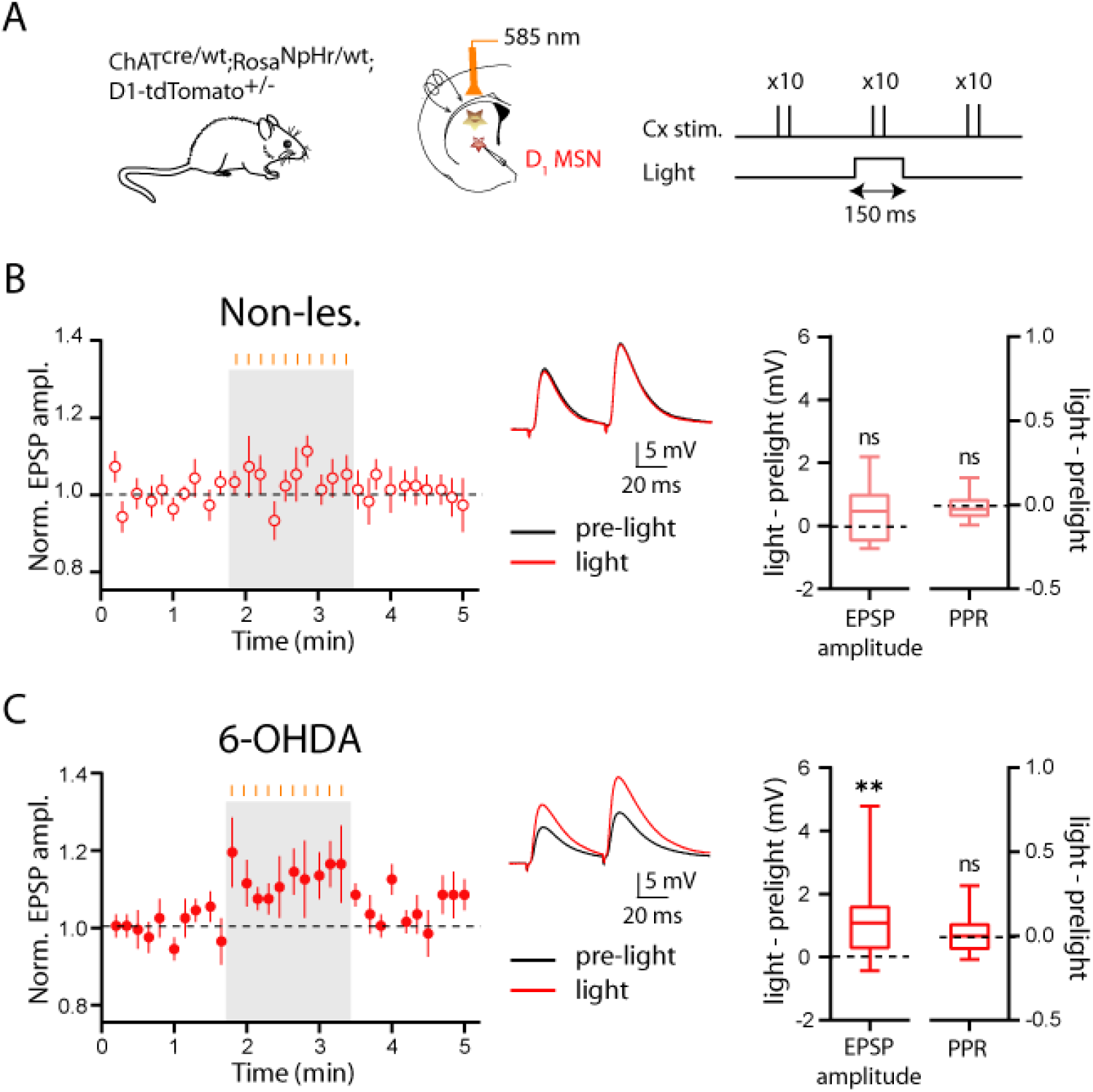
Corticostriatal EPSPs in D1 MSNs are increased by CIN opto-inhibition in 6-OHDA mice. (A) Schematic of the experimental approach. (B, C) Time courses (mean ± SEM) of normalized EPSP amplitude recorded in non-lesioned (B, n = 12) and 6-OHDA (C, n = 13) mice before, during and after CIN opto-inhibition. Light delivery (10 pulses, 150-ms width at 1 Hz) is indicated by the orange vertical bars above the grey rectangle. Representative example of averaged EPSP traces (10 consecutive trials) recorded in non-lesioned (B) and 6-OHDA (C) mice before (pre-light) and during (light) opto-inhibition of CINs. Paired cortical stimulations (50 ms interval) were delivered every 10 s. Box-and-whisker plots illustrate the difference in EPSP amplitude and PPR evoked in light vs. pre-light conditions. Box-and-whisker plots indicate median, first and third quartile, min and max values. ** p < 0.01; ns, not significant. See Table S1 for statistical information.

### Interruption of a signaling pathway involving M4 mAChR and PKA mediates the effect of CIN opto-inhibition on corticostriatal transmission

The finding that CIN opto-inhibition enhances cortically-evoked EPSPs in D1 MSNs in 6-OHDA mice without modifying PPR suggests a post-synaptic locus of action. Abnormal post-synaptic regulation of corticostriatal transmission by CINs could result from changes in ACh release, mAChR expression and/or in mAChR-coupled transduction pathways. Although increased cholinergic tone is usually considered as central to the pathophysiology of PD, whether this is due to increased ACh release remains controversial. To reexamine this issue, we used an electrophysiological readout of mAChR activation by endogenous ACh. For that, we virally expressed G-protein-activated potassium channels (GIRK2) in MSNs that do not express this channel endogenously, by injecting an adeno-associated virus (AAV) encoding tdTomato and GIRK2 into the dorsal striatum (Mamaligas and Ford, 2016). After allowing 26-30 days for AAV-GIRK2 expression, application of the muscarinic agonist carbachol (10 μM) evoked an outward current in tdTomato+ MSNs demonstrating that GIRK2 channels efficiently couple to Gαi-linked mAChR (Figure S5). In the absence of mAChR stimulation, tdTomato+ MSNs exhibited spontaneous inhibitory post-synaptic currents (sIPSCs) that were suppressed by tropicamide (1 μM, n= 3), a preferential antagonist of M4 mAChR, indicating that they result from activation of M4 mAChR (Figure S5). Because these sIPSCs result from the activity-dependent vesicular release of ACh (Mamaligas and Ford, 2016), the frequency of these events can be used as a proxy to evaluate ACh release. We therefore compared the frequency of sIPSCs recorded in non-lesioned and 6-OHDA mice and found no significant difference between the two groups, strongly suggesting that ACh release is not altered in parkinsonian mice (Figure 2A and Table S3). We next measured by RT-qPCR the relative abundance of *Chrma1* and *Chrma4* mRNAs encoding for M1 and M4 mAChR subunits in FACS-sorted D1 MSNs from Drd1a-tdTomato mice, 28 to 33 days post-vehicle or 6-OHDA injection (Figure S6A). Taking *Gapdh* as a reference gene because its expression was not affected by the 6-OHDA lesion (Figure S6B and Table S3), the results showed no significant difference between the two conditions (Figure 2B and Table S3). Taken together, these results suggest that the 6-OHDA-induced changes in cholinergic signaling do not occur through an alteration of ACh release or mAChR gene expression.

**Figure 2:**
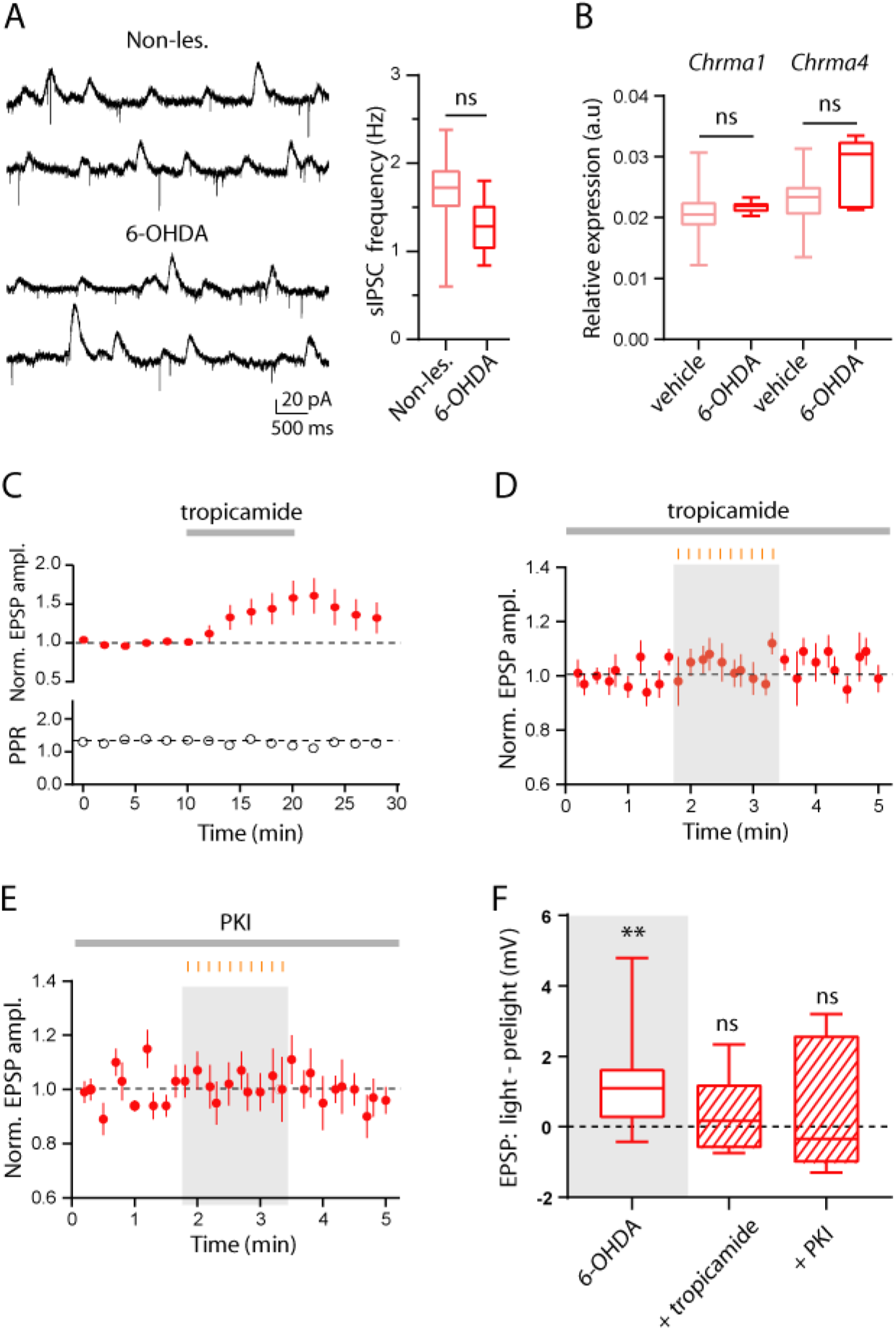
M4 mAChR and protein kinase A mediate the increase of corticostriatal EPSPs induced by CIN opto-inhibition in D1 MSNs in 6-OHDA mice. (A) Representative trace of sIPSCs recorded in tdTomato+ MSNs (Vh = −60 mV) from non-lesioned and 6-OHDA mice. Box-and-whisker plot shows the frequency of sIPSCs in non-lesioned (n = 7) and 6-OHDA (n = 10) mice. (B) Relative expression of *Chrma1* and *Chrma4* mRNA, encoding respectively for M1 and M4 mAChR subunits, in D1 MSNs that were FACS-sorted in vehicle (n = 7) and 6-OHDA (n = 7) mice. (C) Normalized EPSP amplitude (mean ± SEM) as a function of time in D1 MSNs from 6-OHDA mice. 10-min bath-application of tropicamide (1 μM) increased EPSP amplitude without affecting PPR (n = 5). (D, E) Graphs showing normalized EPSP amplitude (mean ± SEM) as a function of time in D1 MSNs from 6-OHDA mice. CIN opto-inhibition was performed in the presence of tropicamide (1 μM, n = 10, D) and the PKA inhibitor, myristoylated PKI 14-22 amide (1 μM, n = 8, E). Light delivery (10 pulses, 150-ms width at 1 Hz) is indicated by the orange vertical bars above the grey rectangle. (F) Box-and-whisker plot illustrate the effects of CIN opto-inhibition on EPSP amplitude in the presence of several pharmacological compounds. For comparison purposes, the effect of CIN opto-inhibition in D1 MSNs from 6-OHDA mice described in figure 1C is also illustrated (grey rectangle). Box-and-whisker plot indicates median, first and third quartile, min and max values. ** p < 0.01; ns, not significant. See also Table S3 for statistical information.

We then asked whether the potentiation of EPSPs induced by CIN opto-inhibition could be due to disruption of a mAChR-linked post-synaptic signaling pathway, focusing on M4 mAChR that is preferentially expressed by D1 MSNs. If M4 mAChR is involved, its pharmacological blockade should not only potentiate corticostriatal transmission, as does CIN opto-inhibition, but also occlude the effect of CIN opto-inhibition. Bath-application of tropicamide (1 μM) produced a marked increase in EPSP amplitude (at 22 min, +61 ± 0.23% of baseline, n = 5) without affecting PPR (Figure 2C). Under tropicamide, CIN opto-inhibition failed to affect corticostriatal EPSPs (Figures 2D, F and Table S3). These results strongly argue for a critical involvement of M4 mAChR in the effect of CIN opto-inhibition on corticostriatal transmission in D1 MSNs of 6-OHDA mice. Activation of Gαi-coupled M4 mAChR inhibits protein kinase A (PKA) activity. Therefore, CINs might affect corticostriatal transmission through post-synaptic modulation of PKA. To test this hypothesis, slices were pre-incubated in ACSF containing a cell-permeable peptide inhibitor of PKA, myristoylated PKI 14-22 amide (1 μM), for at least 1 hour prior to recording and was kept into the bath. In this condition, EPSP amplitude remained unchanged from baseline levels during CIN opto-inhibition (Figures 2E, F and Table S3). Taken together, these results indicate that disruption of a PKA-dependent signaling pathway linked to M4 mAChR mediates the potentiation of corticostriatal synaptic transmission in D1 MSNs from 6-OHDA mice.

### Corticostriatal long-term potentiation is lost *in vivo* in parkinsonian mice and is partially restored by CIN opto-inhibition

Slice experiments have provided compelling evidence for altered long-term plasticity at corticostriatal synapses in PD condition. Whether this also occurs *in vivo* after chronic DA depletion and is related to CIN activity remains an open question. To address this issue, we used a protocol efficient to induce LTP *in vivo* in non-lesioned mice as described in a previous work in rats (Charpier and Deniau, 1997), and determined whether or not this form of plasticity is maintained in 6-OHDA mice. Intracellular recordings of MSNs were performed in the dorsolateral region of the striatum receiving direct inputs from the sensorimotor cortex. MSNs recorded under ketamine-xylazine anesthesia exhibited spontaneous membrane fluctuations consisting of recurrent sustained depolarization (up states) interrupted by hyperpolarizing periods (down states) (Figure 3A). EPSPs were triggered by electrical stimulation of the sensorimotor cortex during the down state to minimize EPSP amplitude fluctuations due to the different level of membrane polarization. To induce LTP, we applied 4 trains of 1 s cortical stimulation at 100 Hz at 10 s interval coupled to membrane depolarization of MSNs (Figure 3A). In experiments where CIN opto-inhibition was coupled to the pairing protocol, light was applied for 1 s at the same time as the trains. As a prerequisite, we verified that CIN firing was reliably inhibited for the duration of the illumination (Figure S7). On average, EPSP amplitude recorded in MSNs of non-lesioned mice was significantly increased after the pairing protocol (130.7 ± 12.7% of baseline, 20-30 min post-pairing, n = 6) (Figure 3B and Table S4). In 6-OHDA mice, the pairing protocol no longer triggered LTP (93.7 ± 7.7% of baseline, 20-30 min post-pairing, n = 5). When the same protocol was applied in 6-OHDA mice in conjunction with opto-inhibition of CINs, the group average also showed no significant LTP (113.1 ± 18.3 % of baseline, 20-30 min post-pairing, n = 5). However, given the high degree of variability of the individual cell responses within the different experimental groups, we compared pre- and post-pairing values in each cell for all conditions (Figure 3C and Table S4). Upon the 6 cells tested in non-lesioned mice, 4 displayed LTP and 2 no plasticity. In 6-OHDA mice, none of the 5 cells recorded showed LTP, 3/5 no plasticity and 2/5 LTD. Finally, in 6-OHDA mice in which CIN opto-inhibition was coupled to the pairing protocol, 2/5 cells displayed LTP, 2/5 no plasticity and 1/5 LTD. Overall, these results show that DA depletion induces a complete loss of LTP *in vivo* that is attenuated by CIN opto-inhibition.

**Figure 3:**
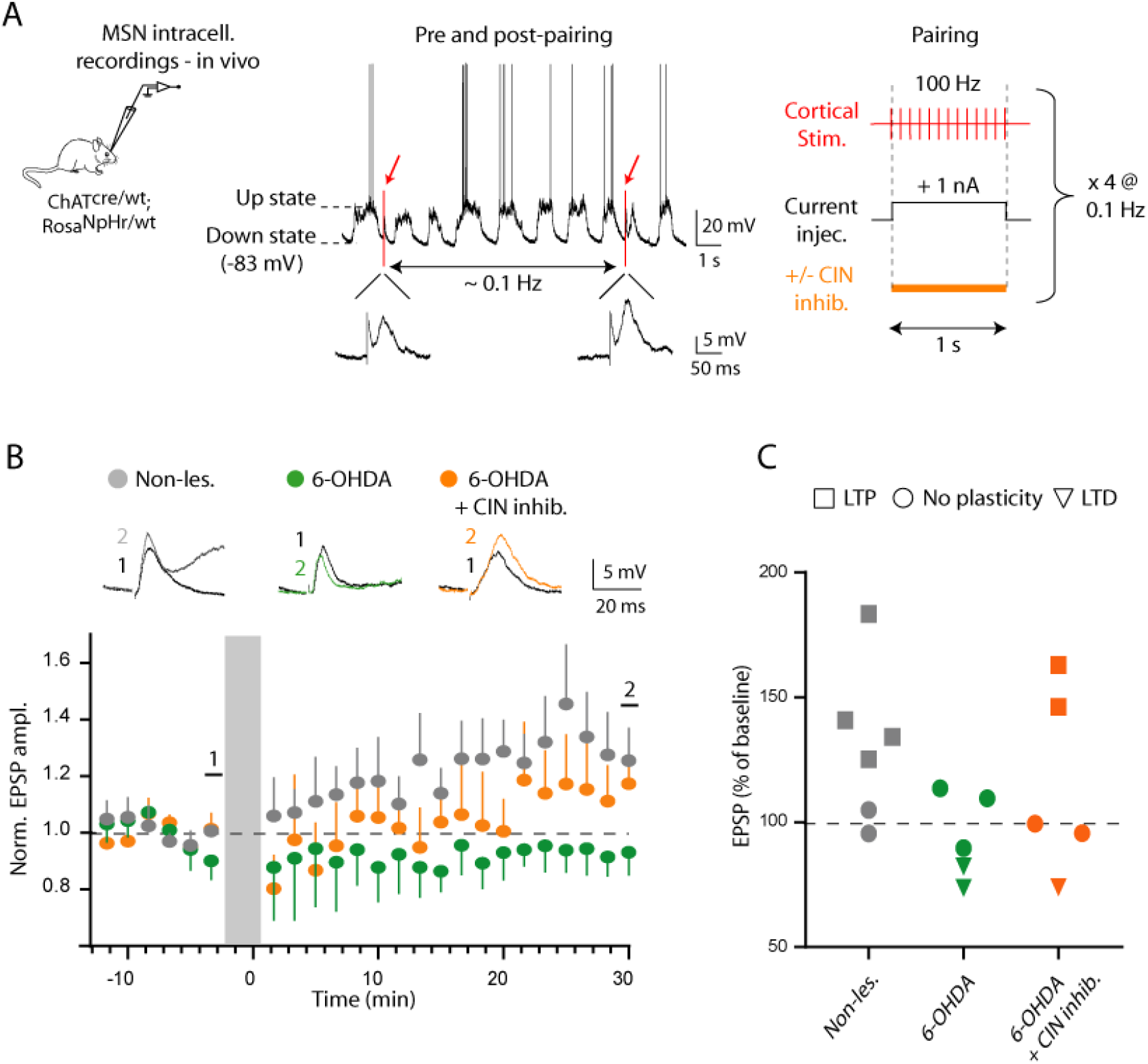
The loss of corticostriatal long-term potentiation *in vivo* in parkinsonian mice is partially restored by CIN opto-inhibition. (A) Example trace of a current clamp recording from one MSN exhibiting up and down membrane fluctuations. Red arrows indicate the time when cortical stimulations were applied during the down states. Expanded traces show the two cortically-evoked EPSPs. Schematic of the experimental protocol used to induce LTP: cortical tetanic stimulation (1 s at 100 Hz applied four times at 10-s intervals) coupled to post-synaptic membrane depolarization (+ 1 nA). The orange bar represents light application during the pairing protocol. (B) Time courses (mean ± SEM) of normalized EPSP amplitude from MSNs recorded in non-lesioned (n = 6), 6-OHDA (n = 5) and 6-OHDA + CIN opto-inhibition (n = 5) conditions before and after the pairing protocol. No recording is performed during application of the plasticity protocol (grey box). Insets show superimposed averaged EPSPs (n=8) for individual cells recorded just before (black traces, 1) and 30 min after application of the pairing protocol (color of experimental group, 2). (C) Summary graph showing the percentage of change in EPSP amplitude 20-30 min after the pairing protocol. Each symbol represents an individual cell and indicates the direction of plasticity change (square: LTP; circle: no plasticity, triangle: LTD).

### Decreasing CIN activity improves motor-skill learning in parkinsonian mice

Altered transmission and plasticity at corticostriatal synapses are considered as key substrates of motor deficits and motor learning impairment in PD state. Because CIN silencing increases corticostriatal transmission onto D1 MSNs, which are hypoactive in PD state, and restores LTP in some MSNs, we hypothesized that silencing of these interneurons might improve motor-skill learning in parkinsonian mice. To test this assumption, we used a chemogenetic approach, more adapted for durable modulation of neuronal activity compatible with behavioral testing. An inhibitory DREADD was selectively expressed in CINs by injecting a Cre-inducible AAV carrying the gene encoding hM4Di fused to mCherry into the striatum of ChAT Cre mice. First, the kinetic profile of CNO-mediated modulation of CIN activity was determined *in vivo* by performing CIN extracellular recordings in anaesthetized hM4Di-injected mice (Figure 4A). Putative CINs were identified by their typical tonic discharges (3.3 ± 1.4 Hz, n = 4) and long-lasting action potentials (> 2.5 ms). CIN firing rate was reduced by ~75% 45 min after i.p. injection of CNO (1 mg/kg) and this decrease in activity remained stable for at least another 30 min (Figure 4B). We then examined the impact of CIN chemogenetic silencing in 6-OHDA hM4Di-injected mice subjected to the rotarod test. Their performance was compared to that of 6-OHDA and non-lesioned mice that received a control AAV carrying only the reporter gene mCherry. All experimental groups received CNO (1 mg/kg, i.p.) 45 min prior to behavioral assessment, so that any potential off-target effects of CNO would affect all mice equally (Figure 4C). Since 6-OHDA mice performed poorly in this test, all mice were first trained for 5 days to stay on the rotating cylinder when it was spinning at a constant speed (12 rpm, cut-off time 60 s). After the training period, we applied an accelerating protocol for 3 days in which the cylinder speed gradually increases from 4 to 40 rpm in 5 min, which is commonly used to evaluate motor-skill learning in rodents (Giordano et al., 2018; Yin et al., 2009). As previously reported in PD mouse models (Giordano et al., 2018), 6-OHDA mice showed impaired motor-skill learning: while the latency to fall off the rod progressively increased in non-lesioned mice, 6-OHDA mice showed no progress during this period (Figure 4D and Table S5). Chemogenetic inhibition of CINs greatly improved the performance as evidenced by the significant increase in the latency to fall off the cylinder across days in 6-OHDA mice injected with the inhibitory DREADD (Figure 4D and Table S5). These data indicate that reducing CIN activity is an effective way to rescue motor-skill learning in parkinsonian mice.

**Figure 4:**
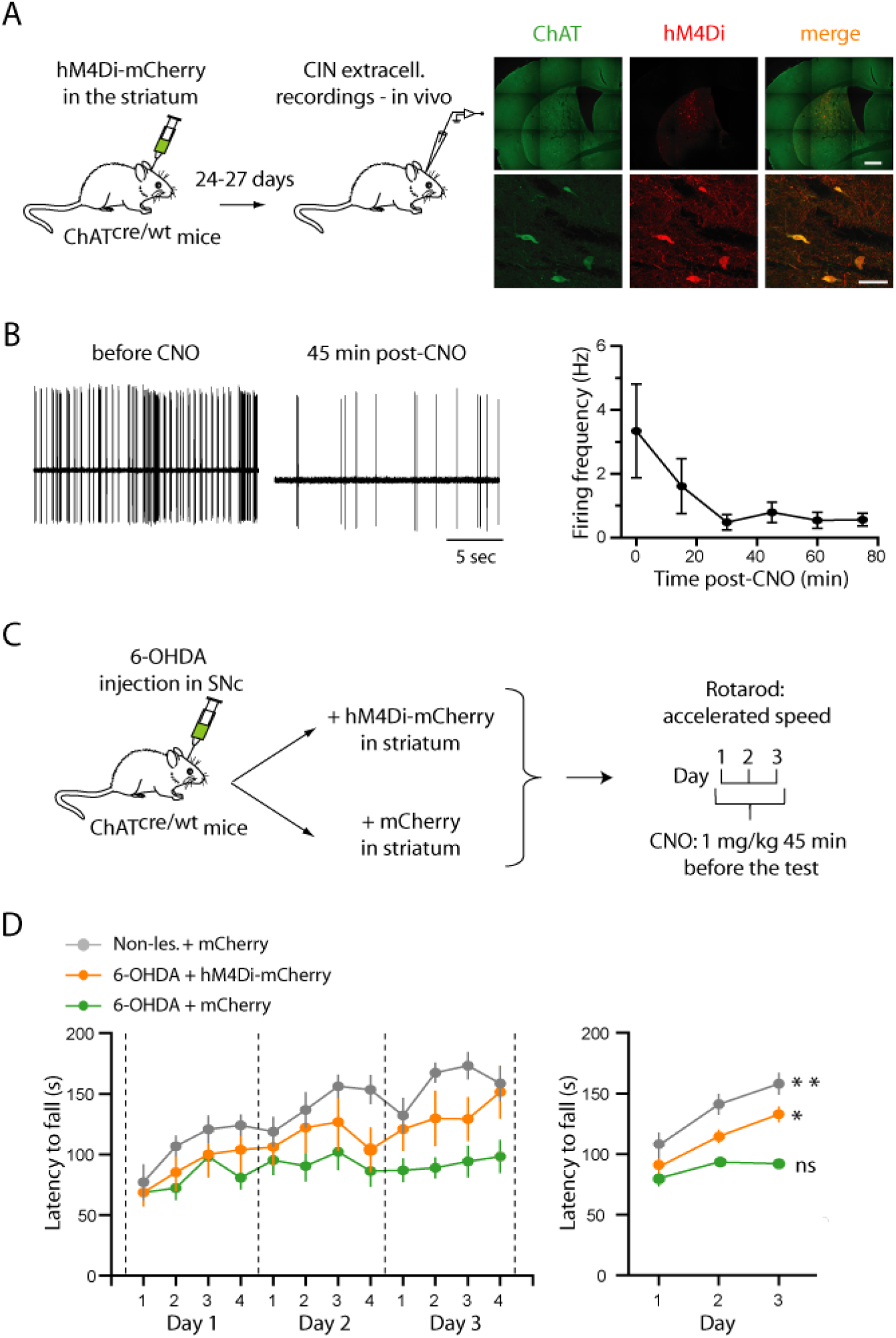
Chemogenetic inhibition of CINs improves motor-skill learning in 6-OHDA mice. (A) Schematic of *in vivo* recordings in anaesthetized mice expressing hM4Di-mCherry and photomicrographs showing the co-localization of DREADD (mCherry, red) with CINs (eGFP, green) in the striatum. Scale bars: top 500 μm, bottom 50 μm. (B) Representative example of one putative-CIN unit recorded *in vivo* and graph (mean ± SEM) showing the spike frequency of putative CINs before and at several time points after i.p. injection of CNO (1 mg/kg, n = 4). (C) Schematic of the experimental design for the rotarod test. (D) Latency to fall off the rotarod (mean ± SEM) across daily trials of each day (left) or averaging trials values (right) in each experimental group (grey: non-lesioned mice injected with AAV-mCherry, n = 9; orange: 6-OHDA mice injected with AAV-hM4Di-mCherry n = 7; green: 6-OHDA mice injected with AAV-mCherry, n = 14). ** p < 0.01; * p < 0.05; ns, not significant. Performances were compared at Day 3 vs. Day 1. See also Table S5 for statistical information.

## DISCUSSION

Modulation of corticostriatal transmission by ACh has been inferred primarily from the action of cholinergic receptor agonists in slices experiments. Here, we used an optogenetic approach to probe the effects of endogenous ACh on corticostriatal transmission, focusing on D1 MSNs. A hallmark of CINs is their continuous tonic activity (Bennett and Wilson, 1999; Bennett et al., 2000) and the stereotypical pauses they acquire in response to salient or reward prediction-related stimuli during conditioning (Apicella et al., 1991). Because these features suggest that a brief interruption in CIN firing is a meaningful signal within the striatum, we opted for inhibitory optogenetic tools to manipulate their activity. Our findings provide the first evidence that acute inhibition of CIN firing potentiates corticostriatal transmission in D1 MSNs in PD-like condition but not in control condition. This potentiation is not due to altered ACh release or mAChR expression but to the interruption of a post-synaptic pathway involving M4 mAChR and PKA. Furthermore, decreasing CIN firing *in vivo* partially restored LTP at corticostriatal synapses and alleviated motor-skill learning deficits in parkinsonian mice. Overall, these data provide novel mechanistic insight onto the positive effect of anticholinergic strategies in PD.

Previous slice recordings showed that activation of mAChR reduces excitatory transmission onto MSNs (Ding et al., 2010; Hernandez-Echeagaray et al., 1998; Malenka and Kocsis, 1988; Pancani et al., 2014) and, conversely, one study reported that pharmacological blockade of mAChR modestly increases the amplitude of glutamatergic responses in MSNs (Pakhotin and Bracci, 2007). Because most of these manipulations triggered an alteration in PPR, these effects have been attributed to the activation of pre-synaptic mAChR expressed by cortical terminals (Hersch et al., 1994). Consistently, we also observed a pre-synaptic cholinergic inhibition of corticostriatal transmission when ACh level was increased by an acetylcholinesterase inhibitor, neostigmine. However, we found that reducing cholinergic tone via inhibition of CIN firing does not affect cortically-evoked EPSPs and PPR in D1 MSNs from non-lesioned mice. This is consistent with the lack of clear effects on behavior and MSN activity reported in several studies upon CIN opto-inhibition (English et al., 2011; Maurice et al., 2015; Ztaou et al., 2018 but see Zucca et al., 2018). The lack of modulation of corticostriatal transmission is not due to insufficient cholinergic tone in slices as opto-inhibition of CINs also failed to modulate EPSPs in vivo and our recordings of sIPSCs in GIRK2-overexpressing MSNs showed that phasic activation of post-synaptic mAChR occurred in slices. Therefore, our finding strongly suggests that ACh does not exert a tonic control onto corticostriatal transmission at either pre- or post-synaptic levels in non-lesioned mice, which does not exclude phasic activity-dependent modulation in specific behavioral context. In sharp contrast, we found that brief inhibition of CIN firing significantly increased corticostriatal transmission in parkinsonian condition and we provided evidence for a post-synaptic site of action. For instance, there was no change in PPR during CIN opto-inhibition and the preferential antagonist of M4 mAChR, which occluded the potentiation induced by CIN opto-inhibition, substantially increased the amplitude of EPSPs without affecting PPR. These results are fully consistent with a prominent role of post-synaptic M4 mAChR expressed by D1 MSNs. Although we cannot rule out a contribution of brainstem cholinergic afferences, previous results showing that these inputs did not modulate DA release in the striatum whereas targeted activation of CINs did strongly suggest that the action of ACh is, at least under certain conditions, primarily mediated by CINs (Brimblecombe et al., 2018; Dautan et al., 2014). We found that EPSP potentiation occurred via a PKA-dependent mechanism. In the striatum, PKA phosphorylation of the GluA1 subunit is increased after pharmacological blockade of M4 mAChR and results in enhanced AMPAR-mediated currents (Mao et al., 2018; Yan et al., 1999). It is quite likely that CIN inhibition, by decreasing M4 mAChR stimulation, leads to a similar scenario. A puzzling question is why this phenomenon is only observed after DA lesion? The most obvious consequence of DA lesion for D1 MSNs is the loss of stimulation of the D1R-mediated signaling pathway, which increases cyclic adenosine monophosphate (cAMP) levels and activates PKA via Gαs/olf proteins, both *in vitro* (Nair et al., 2019) and *in vivo* (Lee et al., 2021). Therefore, because the levels of ACh are elevated relative to DA in the parkinsonian state (McKinley et al., 2019), the system is biased toward activation of Gαi-coupled M4 mAChR, likely resulting in low PKA activity (Lerner and Kreitzer, 2011). In this context, the release of PKA inhibition triggered by CIN silencing would be sufficient to increase AMPAR-mediated responses specifically in parkinsonian mice. Similar disinhibition in healthy condition would be lessened by the stimulatory effect of D1R and would not be potent enough to modulate MSN glutamatergic responses, at least under basal conditions.

In contrast to the long-standing assumption that ACh tone increases as DA level falls, we did not find evidence for a significant change in ACh release in parkinsonian mice. Measurement of ACh concentration in brain samples has been particularly challenging due its rapid hydrolyzation by acetylcholinesterase, likely explaining the conflicting results obtained with this technique. Similarly, the increase in spontaneous CIN activity that would be expected with a hypercholinergic state has not been clearly established in PD models (Tubert and Murer, 2021). However, other mechanisms such as aberrant synchronization (Raz et al., 2001) or alteration of RGS4-dependent auto-receptors (Ding et al., 2006) may explain the increased impact of cholinergic transmission in PD. Here, the use of a direct electrophysiological readout of mAChR activation allowed us for the first time to probe ACh release in non-lesioned and 6-OHDA mice and our results did not reveal major alterations of ACh release under parkinsonian conditions. Finally, DA lesion did not affect either the expression of M1 and M4 mAChR in D1 MSNs, reminiscent of the results obtained for nicotinic receptors expressed by thalamic afferences (Tanimura et al., 2019). Overall, given the lack of significant changes in ACh release and mAChR expression in the parkinsonian state, an imbalance between cholinergic signaling via M4 mAChR and dopaminergic signaling via D1R is probably the most prominent alteration induced by DA lesion in D1 MSNs. This imbalance could result in tonic inhibition of corticostriatal transmission in these neurons by CINs, explaining why their optogenetic silencing or the pharmacological blockade of the M4 mAChR potentiates corticostriatal transmission. CIN-mediated tonic inhibition could account for the reduction in the strength of corticostriatal synapses observed in D1 MSNs after DA lesion (Fieblinger et al., 2014). Interestingly, a previous work demonstrated that the loss of D2R signaling contributed to the amplification by CINs of thalamic-evoked responses in D2 MSNs in parkinsonian mice (Tanimura et al., 2019). Together, these data suggest that suppression of DA action on D1R and D2R triggers an abnormal control of corticostriatal and thalamostriatal transmission by CINs in parkinsonian state.

Long-term changes in the synaptic strength of corticostriatal synapses are believed to represent the cellular substrate of sensorimotor learning (Di Filippo et al., 2009; Graybiel, 1998; Pisani et al., 2005). More precisely, the dorsolateral striatum that receives projections from somatosensory and motor cortical areas has been implicated in the acquisition and performance of motor sequences and could be activated during all stages of incremental motor learning (Giordano et al., 2018; Thorn et al., 2010 but see Yin et al., 2009). We therefore recorded MSNs in the dorsolateral area of the striatum and found that a Hebbian protocol induced LTP of corticostriatal transmission in a majority of MSNs in non-lesioned mice, in agreement with earlier works (Charpier and Deniau, 1997; Fisher et al., 2017). In contrast, MSNs from parkinsonian mice no longer exhibited LTP. This is, to the best of our knowledge, the first report showing the loss of this form of plasticity *in vivo* after chronic DA lesion and this result is consistent with *in vitro* data (Centonze et al., 1999; Picconi et al., 2003). In this condition, MSNs showed no net plasticity, some even displaying LTD, which corroborates previous results obtained both *in vivo* after acute DA depletion (Reynolds and Wickens, 2000) and in slices experiments (Shen et al., 2008). More precisely, in this latter work, the protocol that normally leads to LTP in D1 MSNs induces LTD after DA depletion, suggesting that corticostriatal plasticity may be affected in a cell-specific manner and is dysfunctional rather than absent in PD.

We and others have previously shown that targeted inhibition of CINs improved performance of parkinsonian mice in motor tasks that do not have a strong learning component (Maurice et al., 2015; Tanimura et al., 2019; Ztaou et al., 2016). Our present result demonstrating that chemogenetically-induced decrease in CIN firing enabled parkinsonian mice to significantly improve their performance over time in the accelerating rotarod task extends this notion to motor-skill learning. Considering that corticostriatal LTP is induced by motor learning in the dorsolateral striatum (Giordano et al., 2018), our working hypothesis is that CIN inhibition improves motor learning in parkinsonian mice through their ability to induce LTP at corticostriatal synapses, as observed here in some MSNs. We showed that decreasing cholinergic tone on M4 mAChR potentiates basal corticostriatal transmission via a PKA-dependent mechanism in D1 MSNs. Interestingly, PKA is also activated in these neurons in the early learning stages (Lee et al., 2021) and this activation could promote PKA-dependent LTP (Picconi et al., 2003). LTP induction is instead abolished in D1 MSNs when M4 mAChR signaling is boosted with positive allosteric modulator (Shen et al., 2015). These results suggest that inhibition of CIN firing in parkinsonian state might favor PKA-dependent LTP in D1 MSNs by lowering M4 mAChR activation.

While further studies are needed to fully understand how CINs impact the striatal network in normal and pathological conditions, our data reveal the existence of abnormal ACh responsiveness in D1 MSNs in parkinsonian state that impacts corticostriatal transmission. They also suggest that targeting CIN firing is a valuable therapeutic option that avoids extrastriatal and unspecific effects of anticholinergic drugs (Paz and Murer, 2021).

## ACKNOWLEDGMENTS

This work was funded by grants from Fondation de France and Association France Parkinson. It was supported by the “Centre National de la Recherche Scientifique” (CNRS), Aix-Marseille University (AMU) and the French government under the Programme “Investissements d’Avenir", Initiative d’Excellence d’Aix-Marseille Université via A*Midex funding (NeuroMarseille, AMX-19-IET-004) and ANR (ANR-17-EURE-0029). JP was supported by a PhD fellowship from the Ministry of Higher Education, Research and Innovation. We thank Jean-Pierre Kessler for statistical analysis, Maxime Assous, Eduardo Gascon, Aziz Moqrich and Hélène Marie for their critical comments on the manuscript.

## AUTHOR CONTRIBUTIONS

A.R., L.K-LG., N.M., C.B. designed the study. G.L., J.P., A.R., C.M., N.M., C.B. performed experiments. G.L., J.P., A.R., N.M., C.B. analyzed the results. N.M. and C.B. contributed to funding acquisition. C.B. wrote the manuscript with the participation of N.M. and L.K-LG. All authors read and approved the submission.

## DECLARATION OF INTERESTS

The authors declare no competing financial interests.

---

**Figure S1:**
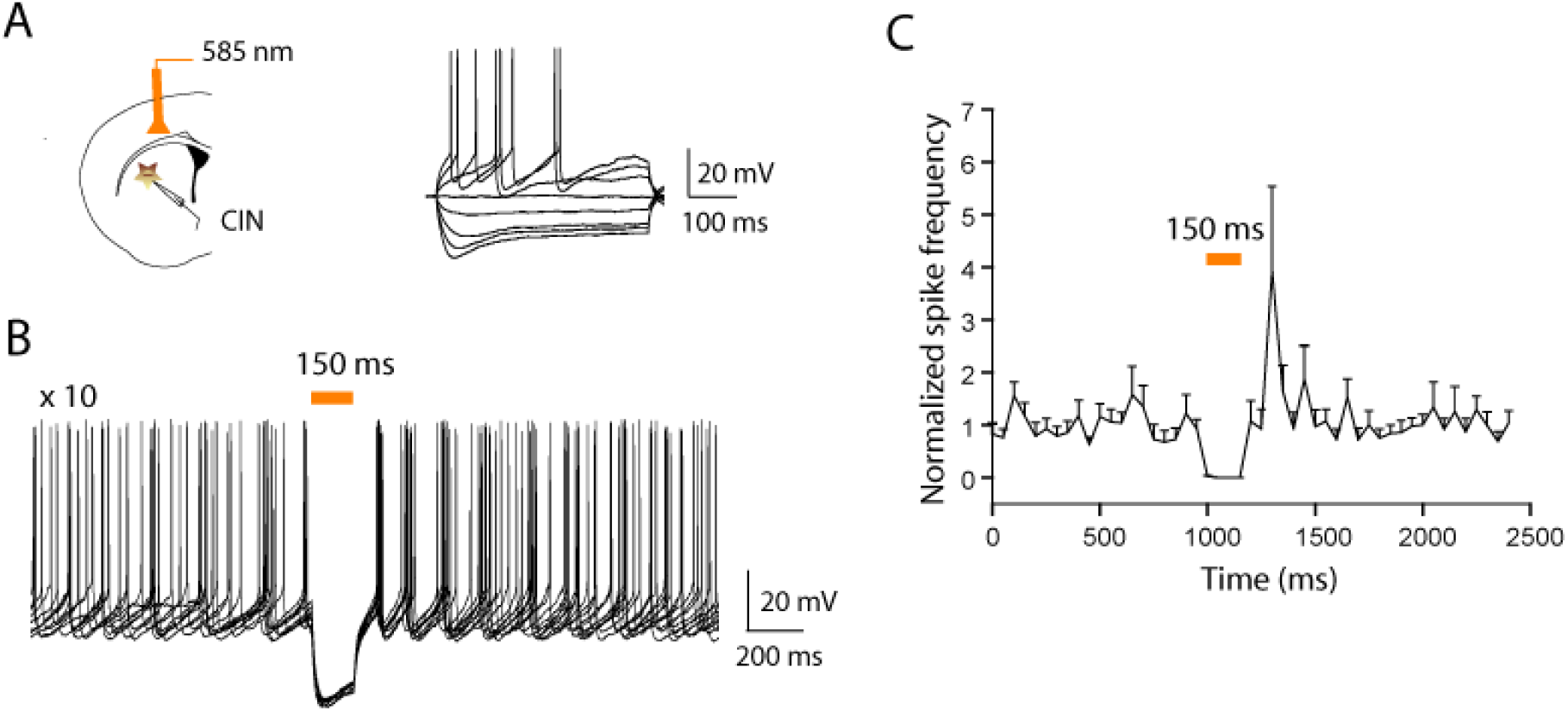
The spontaneous firing of eNpHR-expressing CINs is consistently inhibited by short-duration pulses of 585 nm light *in vitro*. (A) Schematic of CIN activity recordings in slices from transgenic mice expressing eNpHR in CINs. Membrane potential changes induced by current injection in one eNpHR- expressing CIN (−200 to +150 pA, 50 pA increment). (B) Current-clamp recording of the spontaneous firing of an individual CIN subjected to 150 ms light illumination repeated 10 times. (C) Normalized firing frequency (mean ± SEM) before, during and after 150 ms light application (n = 9).

**Figure S2:**
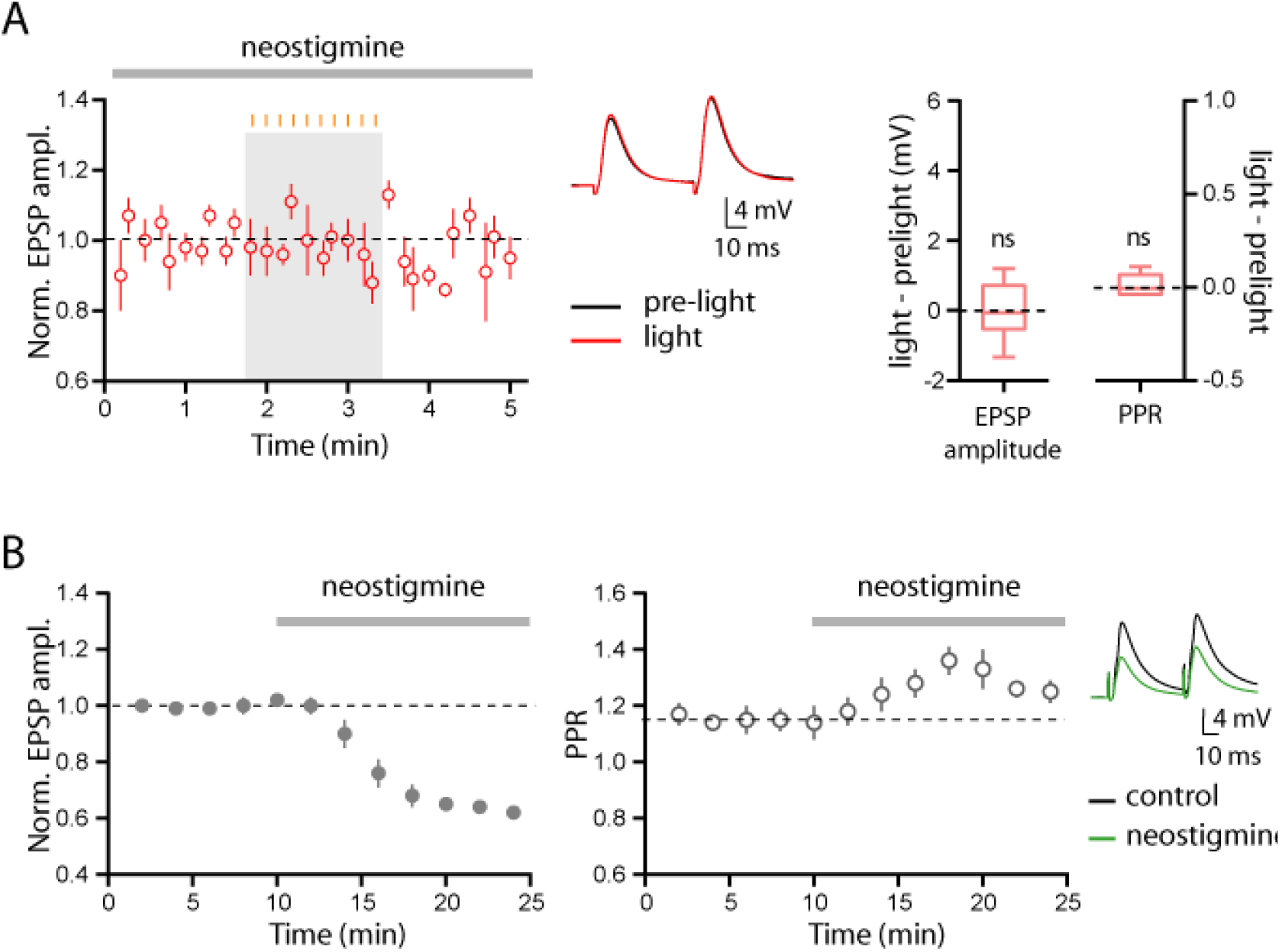
Increasing cholinergic tone does not reveal an effect of CIN opto-inhibition in non-lesioned mice. (A) Time courses (mean ± SEM) of normalized EPSP amplitude in D1 MSNs (n = 6) before, during and after CIN opto-inhibition applied in the presence of neostigmine (3 μM). Light delivery (10 pulses, 150-ms width at 1 Hz) is indicated by the orange vertical bars above the grey rectangle. Representative example of averaged EPSP traces recorded before (pre-light) and during (light) opto-inhibition of CINs in the presence of neostigmine (3 μM). EPSPs are averages of ten consecutive trials. Box-and-whisker plots illustrate the difference in EPSP amplitude and PPR evoked in light vs. pre-light conditions and indicate median, first and third quartile, min and max values. ns, not significant. See Table S2 for statistical information. (B) Time courses (mean ± SEM) of normalized EPSP amplitude and paired-pulse ratio in MSNs (n = 8 cells, N = 4 mice) showing the effect of neostigmine (3 μM, 15 min). Representative example of averaged EPSP traces (ten consecutive traces) before and after bath-application of neostigmine.

**Figure S3:**
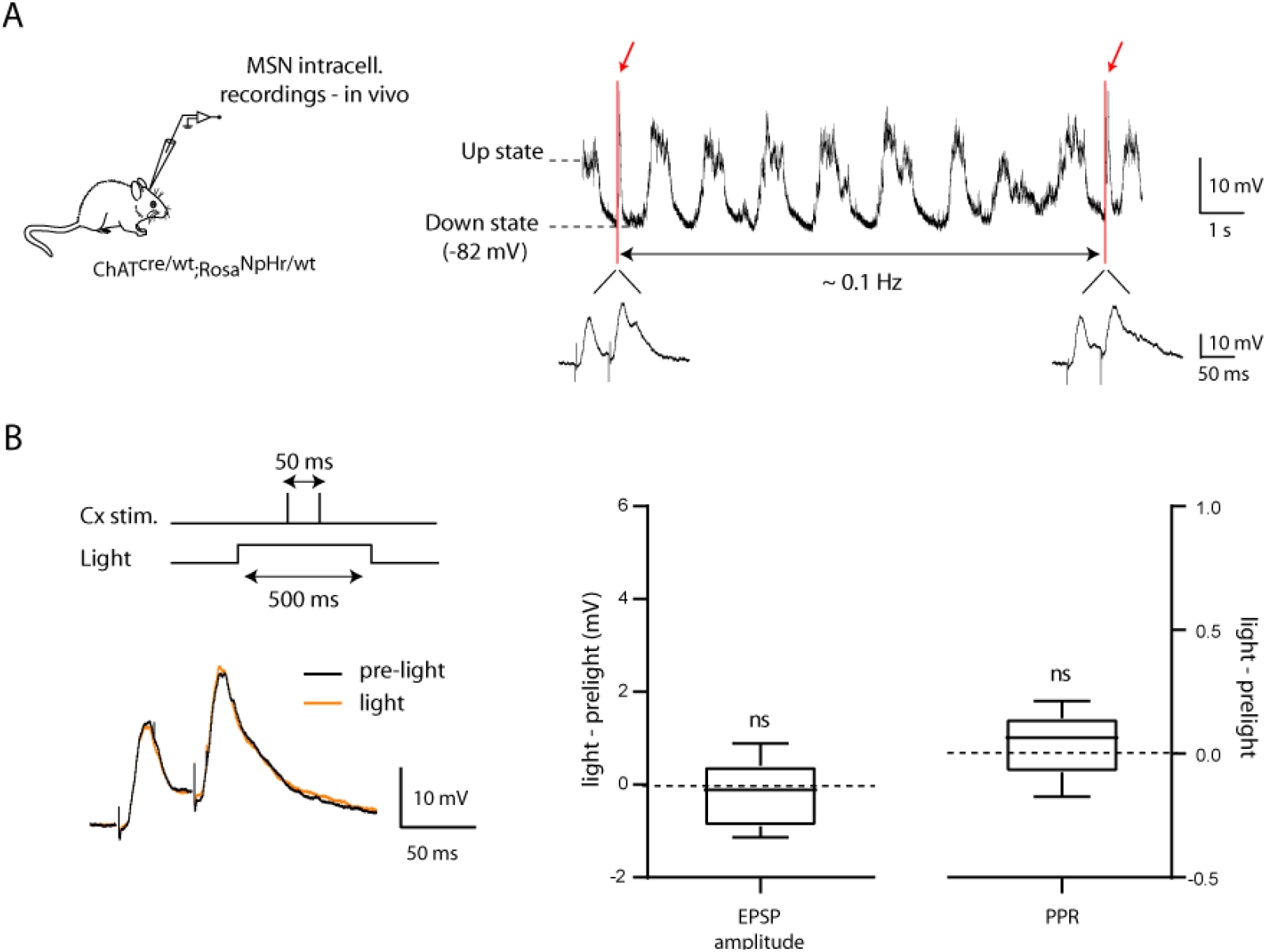
Corticostriatal transmission recorded *in vivo* in MSNs from non-lesioned mice are not affected by CIN opto-inhibition. (A) Example of an intracellular recording from one MSN exhibiting up and down membrane fluctuations. Red arrows indicate the time when cortical stimulations were applied during the down states (paired stimulations at 50 ms interval). Expanded traces show the cortically-evoked EPSPs. (B) Representative example of averaged EPSP traces (8 consecutive trials) in MSNs recorded from non-lesioned mice before (pre-light) and during (light, 500-ms widht) opto-inhibition of CINs. The graphs illustrate the difference in EPSP amplitude and PPR evoked in light vs. pre-light conditions (n = 10). Box-and-whisker plots indicate median, first and third quartile, min and max values. ns, not significant. See also Table S2 for statistical information.

**Figure S4:**
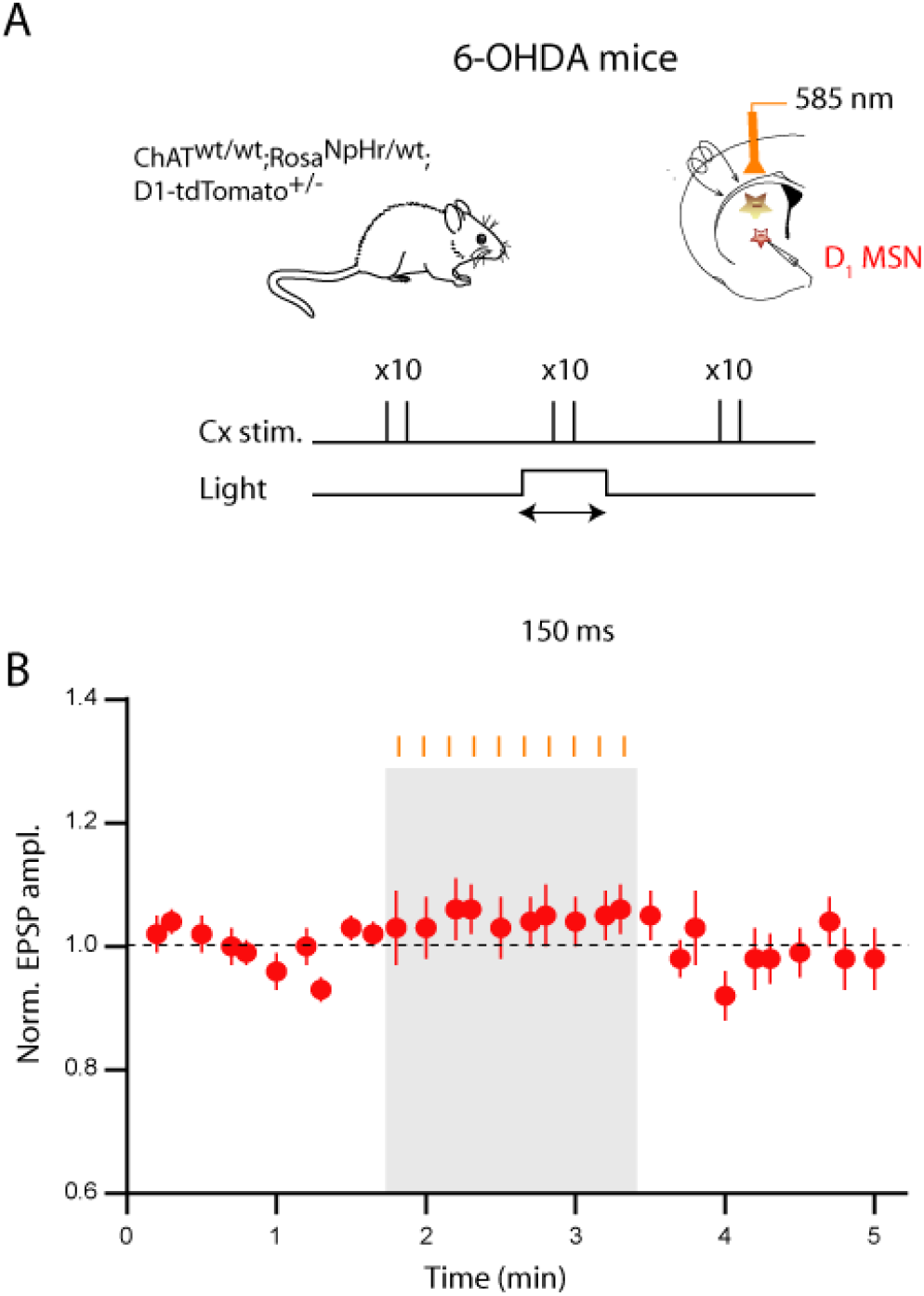
Light application does not modulate EPSPs in 6-OHDA mice that do not express eNpHR. (A) Schematic of experimental protocol. Mice do not express Cre recombinase in cholinergic neurons. (B) Time courses (mean ± SEM) of normalized EPSP amplitude recorded in 6-OHDA mice (n = 18) before, during and after CIN opto-inhibition. Light delivery (10 pulses, 150-ms widht at 1 Hz) is indicated by the orange vertical bars above the grey rectangle. See also Table S2 for statistical information.

**Figure S5:**
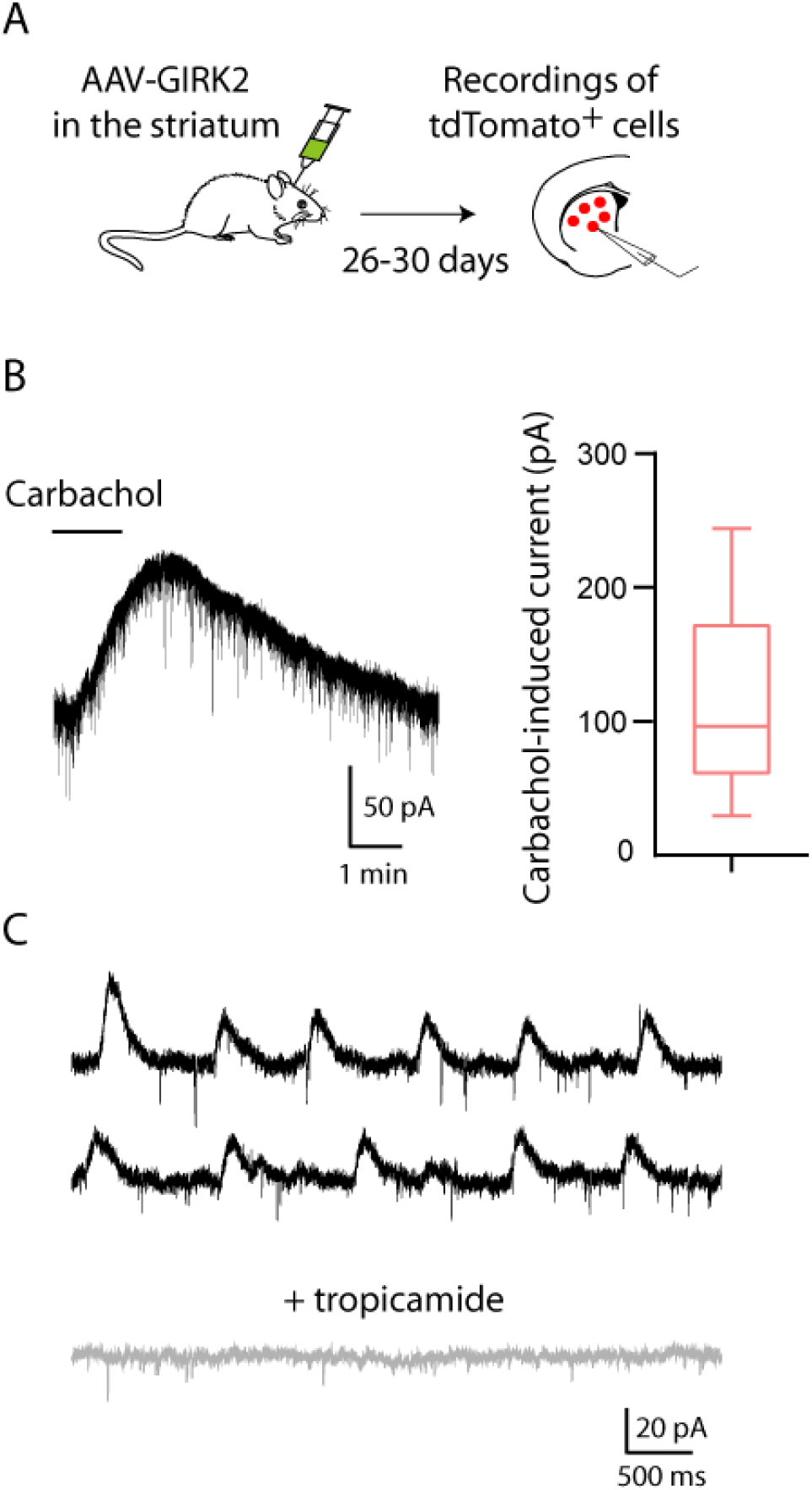
GIRK2 channels couple efficiently to Gαi-linked M4 mAChR. (A) Schematic of the experimental approach. (B) Bath-application of the muscarinic agonist carbachol (10 μM, 2 min) evoked an outward current in a tdTomato+ MSN recorded in voltage-clamp mode (Vh = −70 mV). Box-and-whisker plot shows the amplitude of the carbachol-induced currents in tdTomato+ cells (n = 5). (C) Representative traces illustrating the blockade of sIPSCs recorded in a tdTomato+ MSN (Vh = −60 mV) by the preferential antagonist of M4 mAChR, tropicamide (1 μM).

**Figure S6:**
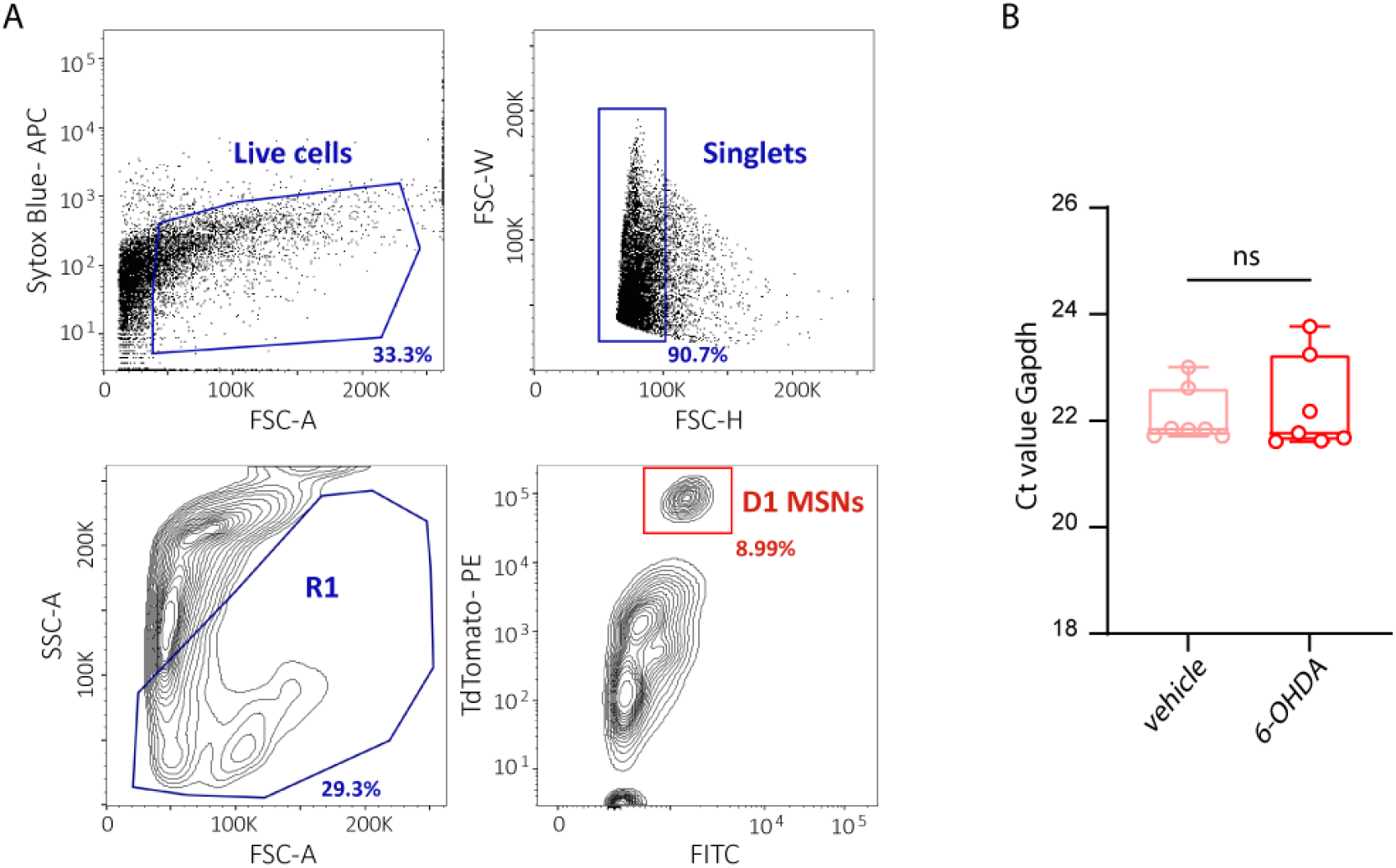
Gating strategy for FACS-sorting of D1 MSNs. (A) Live cells in appropriate size range (FSC-A) were gated by excluding Sytox Blue+ cells, then single cells were gated by excluding doublets in height (FSC-H) and width (FSC-W) dimensions. R1 gate was further drawn according to pre-determined size (FSC-A) and structure (SSC-A) parameters and tdTomato+ D1 MSNs were sorted from R1. Numbers indicate the percentage of events in the corresponding gate. The graphs show one representative experiment out of 14 (7 for vehicle-injected and 7 for 6-OHDA-injected mice). (B) Ct values for *Gapdh* for each experimental group. See also Table S3 for statistical information.

**Figure S7:**
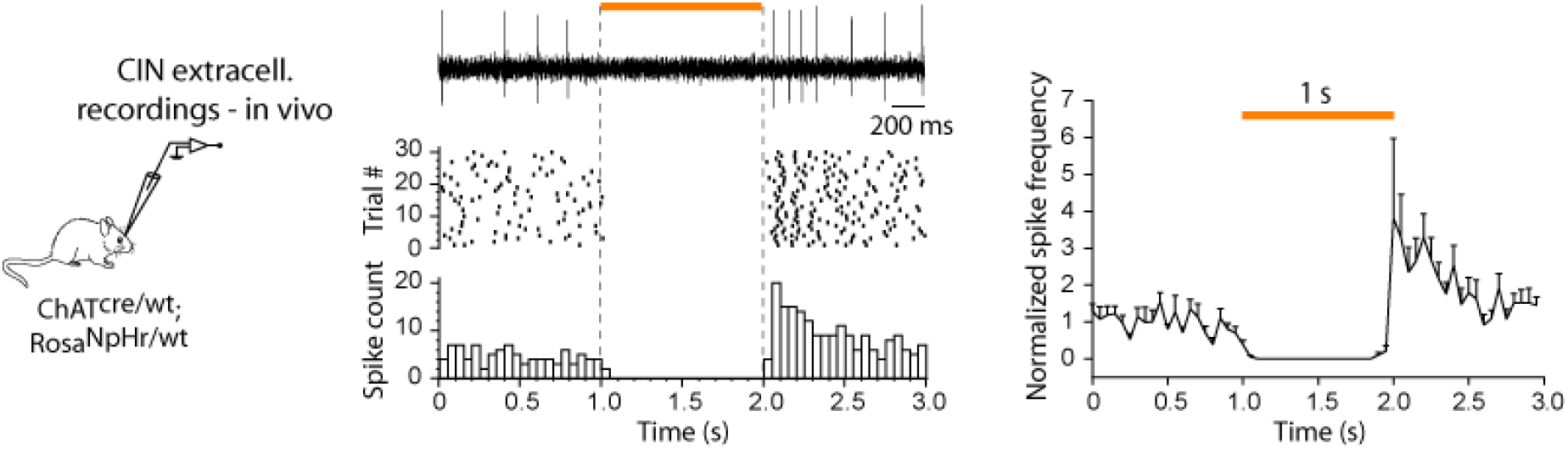
The spontaneous firing of eNpHR-expressing CINs is consistently inhibited by 1-second light pulses *in vivo*. Left, Raster and PSTH obtained in one eNPHR-expressing CIN recorded extracellularly *in vivo* in anaesthetized mice showing the effect of 1sec-duration pulses of 585 nm light. Right, normalized firing frequency (mean ± SEM) of eNpHR-expressing CINs before, during and after 1 s light application (n = 7).

## METHODS

### Mice

All mice strains used in this study were purchased from Jackson Laboratory. We used Choline acetyltransferase ChAT-IRES-Cre knock-in mice (stock number: 006410) and LoxP-stop-eNpHR3.0-EYFP mice (stock number: 014539). Homozygotes from each strain were crossed in-house to produce heterozygous, double-transgenic ChAT^cre/wt^;Rosa^NpHR/wt^ offspring and were used for *in vivo* electrophysiology. For *in vitro* electrophysiological experiments requiring D1 MSN identification and NpHR expression in cholinergic neurons, homozygous ChAT^cre/cre^ mice were first crossed with BAC Drd1a-tdTomato mice (D1-tdTomato mice, stock number: 016204). Double transgenic offspring (ChAT^cre/wt^;D1-tdTomato^+/−^) were then crossed with homozygotes Rosa^NpHR/NpHR^ to produce about one third of triple-transgenic ChAT^cre/wt^;D1-tdTomato^+/−^;Rosa^NpHR/wt^ mice. To provide chemogenetic inhibition of CINs, AAV containing the inhibitory DREADDs was injected into the striatum of heterozygous ChAT^cre/wt^ mice. Both adult male and female mice were used in this study. Animals were housed 3-5 per cage (1-2 per cage if animals underwent surgical procedures), maintained under standard housing conditions (12-hour light/dark cycles, 23°C, 40% humidity) with unrestricted access to food and water and monitored daily by animal care technicians and research staff. All procedures agreed with European Union recommendations for animal experimentation (2010/63/EU) and were approved by the national and local ethical committees (Marseille ethics committee # 14). The project authorization is registered under the number2019022216247615-V3 #19414 delivered by the French Ministry of Higher Education, Research and Innovation.

### Stereotaxic surgery

Mice were anaesthetized with intraperitoneal (i.p.) injections of ketamine and xylazine (100 and 10 mg/kg, respectively) and mounted on a stereotaxic apparatus (Kopf Instruments). Injections were made with a 10-μl syringe, connected to a 33- or 34-gauge needle injector by a polyethylene tubing, and controlled by an injection pump at 0.3 μl/min. All stereotaxic coordinates are calculated relative to bregma. Mice received one unilateral injection of 6-OHDA hydrochloride (1.5 μl at 2.7 μg/μl diluted in 0.9% sterile NaCl containing 0.1% ascorbic acid, Sigma-Aldrich) into the substantia nigra pars compacta (−3.0 mm AP, ±1.3 mm ML, −4.3 mm DV). For cre-dependent hM4Di-mcherry expression in CINs, 6-OHDA injection was immediately followed by unilateral injection of AAV8-hSyn-DIO-hM4D(Gi)-mCherry (2.9×10^13^ GC/mL, Addgene) into the striatum ipsilateral to the lesioned side of ChAT^cre/wt^ mice. Injections were performed at four sites (0.75 μl per site) at the following coordinates: +1.1 mm AP, ±1.7 mm ML, −2.7 and −2.1 mm DV and +0.38 mm AP, ±1.9 mm ML, −2.7 and −2.1 mm DV. The control virus (AAV8-hSyn-DIO-mCherry, 2.3×10^13^ GC/mL, Addgene) was injected according to the same procedure. To express GIRK2 in MSNs, unilateral injection of AAV9-hSyn-tdTomato-GIRK2 (0.5 μl) were performed into the striatum ipsilateral to the 6-OHDA-lesioned side (+0.8 mm AP, ±1.8 mm ML, −2.5 mm DV) (AAV9.hSyn.tdTomato.T2A.mGIRK2-1-A22A.WPRE.bGH, 4.42×10^13^ GC/mL, Penn Vector Core). After injections, the needle was left in place for 10 minutes before removal. The scalp was then sutured and animals were maintained on a heating pad until they fully recovered from anesthesia. Rimadyl (5 mg/kg) was administered subcutaneously for post-operative analgesia over a period of 24-48 hours, including the day of surgery.

### *In vitro* Electrophysiology and optogenetics

Mice were deeply anesthetized with ketamine/xylazine (100/10 mg/kg, i.p.) and transcardially perfused with an ice-cold N-methyl D-glucamine (NMDG)-based solution containing (in mM): 93 NMDG, 2.5 KCl, 1.2 NaH_2_PO_4_, 30 NaHCO_3_, 20 HEPES, 20 glucose, 10 MgCl_2_, 93 HCl, 2 Thiourea, 3 sodium pyruvate, 12 N-acetyl cysteine and 0.5 CaCl_2_ (saturated with 95% O_2_ and 5% CO_2_, pH 7.2-7.4). The brain was then removed from the skull and glued to the stage of a vibratome (Leica, VT1000S) where it remained submerged in ice-cold oxygenated NMDG-based solution. Coronal slices (250 μm thick) containing the striatum were collected. Slices were immediately transferred to recover in NMDG-based solution at 35°C for 5 min and then stored for at least 1h at room temperature in normal artificial cerebrospinal fluid (ACSF) containing (in mM): 126 NaCl, 2.5 KCl, 1.2 MgCl2, 1.2 NaH2PO4, 2.4 CaCl2, 25 NaHCO3 and 11 glucose, to which 250 μM kynurenic acid and 1 mM sodium pyruvate had been added. For the recordings, slices were transferred one at a time to a submersion-type chamber and perfused continuously with warm ACSF (32-34°C) at a rate of 3 ml/min. Solutions were continuously equilibrated with 95% O2 / 5% CO2. In all experiments picrotoxin (50 μM) was included in the ACSF to block GABA_A_ receptors. For PKA inhibition experiments, slices were pre-incubated in ACSF containing PKI 14-22 amide, myristoylated (PKI, 1 μM) for 1.5 hours prior to recording. PKI (1 μM) was kept in the bath during recordings. All compounds were purchased from Tocris or Sigma-Aldrich.

Neurons were visualized on an upright microscope (Nikon Eclipse FN1) equipped with DIC optic and filters set to visualize tdTomato and EYFP using an IR 40x water-immersion objective (Nikon). Combination of electrophysiological properties and expression of fluorophore was used to identify CINs and D1 MSNs. Patch-clamp recordings were performed in whole-cell configurations in current-clamp mode. Patch-clamp electrodes (4-6 MΩ) were prepared from filamented borosilicate glass capillaries (PG150T-7.5, Harvard Apparatus) using a micropipette puller (PC-10, Narishige) and were filled with an intracellular solution containing (in mM): 126 KMeSO4, 14 KCl, 3 MgCl2, 0.5 CaCl2, 5 EGTA, 10 HEPES, 2 NaATP and 0.5mM NaGTP, 10 Na-Phosphocreatine, pH adjusted to 7.25 with NaOH and osmolarity adjusted to 270-280 mOsm/L.

Recordings were obtained using motorized micromanipulators (MP-85, Sutter Instrument), a Multiclamp 700B amplifier, a Digidata 1550B digitizer, and pClamp 10.7 acquisition software (Molecular Devices, San Jose, CA, USA). Signals were low-pass filtered at 10 kHz online and sampled at 10 kHz. Electrode capacitances were compensated electronically during recording. In current-clamp mode, the bridge was continuously balanced and input resistances were monitored with a 50-pA negative step given with every afferent stimulus. Cells showing more than 20% of input resistance variation were excluded from the analysis. To evoke EPSPs in MSNs, electrical stimulation of cortical afferents was performed using a bipolar tungsten electrode placed in the corpus callosum at the border of the cortex and the striatum. As the characteristics of CINs and MSNs can vary according to the territories of the striatum, recordings were restricted to the dorsolateral region of the striatum.

The intensity of the electrical stimulations performed by current pulses (0.1-ms width every 10 s) were adjusted so as to obtain EPSPs with an amplitude approximately equal to 50% of the maximum response. Optogenetic stimulation of eNpHR-expressing CINs was delivered under the control of the acquisition software via the microscope objective lens using wide-field 585 nm LED illumination (Spectra Light Engine, Lumencor, Optoprim). In the majority of experiments examining the effects of CIN photoinhibition on EPSPs, light stimulation was delivered as a 150-ms width pulse. Paired-cortical stimulation (50 ms interstimulus interval) started 50 ms after light was turned on. EPSP amplitudes were calculated by taking the maximum peak amplitude and comparing this with the mean of a 10-ms window immediately before the stimulation artifact. For 6-OHDA injected mice, recordings were performed 20 to 36 days post-injection. For AAV-GIRK2 injected-mice, recordings were performed 26 to 30 days post-injection.

### *In vivo* Electrophysiology and optogenetics

Mice were deeply anesthetized with a mixture of ketamine/xylazine (100/10 mg/kg, i.p. and supplemented as needed during the course of the experiment) and mounted in a stereotaxic head frame (Horsley-Clarke apparatus; Unimécanique, Epinau-sur-Seine, France). Body temperature was maintained at 36.5°C with a homeothermic blanket controlled by a rectal probe (Harvard Apparatus, Holliston, MA). Extracellular recordings: single-unit activity of CINs was recorded extracellularly using glass micropipettes (25-35 MΩ) filled with a 0.5 M sodium chloride solution. Single neuron action potentials were recorded using the active bridge mode of an Axoclamp-2B amplifier (Molecular Devices, San Jose, CA), amplified, and filtered with an AC/DC amplifier (DAM 50; World Precision Instruments). CINs were identified by their classically defined electrophysiological characteristics: large spikes (width > 2.5 ms) and ability to present tonic discharges (> 1.0 Hz). Intracellular recordings: bipolar concentric stainless-steel electrode was inserted in the sensorimotor cortex (A: 2.0 mm; L: +1.5 mm; H: −0.6 mm from the cortical surface). Optical fiber (200 μm-diameter, 0.22 NA; Doric lenses, Québec, CA) was inserted in the striatum with a 15° angle and connected to a 100 mW laser (Combined dual wavelenght, DPSS laser system, Laserglow technologies, Toronto, Ontario, Canada). The entry point had the following coordinates: A: 0.7 mm; L: +2.6 mm. The fiber tip was at a depth of 1.5 mm form the cortical surface. Intracellular recordings of MSNs were performed while simultaneously recording the ECoG using a low-impedance (around 60 kΩ) silver electrode placed on the dura above contralateral motor cortex (A: 2.0 mm; L: −1.5 mm). This allows a precise on-line monitoring of the anesthesia level. Intracellular recordings were performed using 2 M K-acetate-filled glass microelectrodes (40-80 MΩ) and placed in the sensorimotor striatal region (A: 0.7 mm; L: +2.1 mm; H: −1.7/−2.7 mm) related to the stimulated cortical area. All recordings were obtained using an Axoclamp-2B amplifier (Molecular Devices, San Jose, CA) operated in the bridge mode. Data were sampled (300 Hz for ECoG and 25 kHz for intracellular recordings) and stored on-line on a computer connected to a CED interface using the Spike2 data acquisition software (Cambridge Electronic Design, Cambridge, UK). Impaled neurons were considered acceptable when their membrane potential was at least at −60 mV with spontaneous oscillations of large amplitude (>10 mV) and a spike amplitude > 60 mV. Neurons that did not filled these criteria were discarded from the analysis. To measure the input resistance of MSNs, current pulses (200 ms duration applied at 0.5 Hz) were intracellularly injected through the recording electrode. As classically reported for MSNs recorded *in vivo* under ketamine/xylazine anesthesia (Mahon et al., 2001), the cells displayed slow membrane potential fluctuations consisting of recurrent sustained depolarizing plateaus interrupted by hyperpolarizing periods called down states. Through the recording session, cortically evoked EPSPs were induced by triggering the cortical stimulation during the down phase of the membrane potential oscillation using a window discriminator set to detect these down states. Test stimulations (600-μs duration) in the sensorimotor cortex were applied at 0.1 Hz with an intensity below threshold for action potential induction. Two successive cortical stimulations were separated by a 10-s interval plus the time needed (few hundreds ms) for the next down state to occur and trigger the following cortical stimulation. In some cases, the down state was shorter than usual, triggering a cortical stimulation but with an EPSPs evoked after the end of the down state. Such EPSPs were not analyzed. The amplitude of the EPSPs evoked during down state were measured from the baseline to the peak response, then averaged by bins of 100 s (n = 7-10 sweeps) and normalized. Induction of LTP at corticostriatal synapses: a cortical tetanization (100 Hz, 1-s duration repeated 4 times at 0.1 Hz) was delivered in conjunction with a 1-s post-synaptic depolarization achieved by intracellular injection of positive current (+1.0 nA). The intracellular injection of positive current induced a depolarization leading the membrane potential to a suprathreshold level for action potential firing. Current injection started 50 ms before and ended 50 ms after the cortical tetanus. For neurons in which the pairing protocol was combined with opto-inhibition of CINs, the light was switched on during the cortical tetanization, concomitantly to the post-synaptic depolarization. The synaptic responses recorded after the tetanus were normalized to pre-pairing values. Recordings were performed 19 to 27 days post-6-OHDA injection.

### Immunohistochemistry

Immunohistochemical detection of TH and mCherry was performed to control 6-OHDA lesion and DREADDs expression, respectively. To ensure the specificity of DREADDs expression in CINs, immunohistochemical labeling of ChAT was also performed. Animals were deeply anesthetized with a mixture of ketamine/xylazine and then transcardially perfused with an ice-cold solution of paraformaldehyde 4% in PBS. After dissection, brains were post-fixed overnight in the same fixative at 4 °C, cryoprotected in 30% sucrose dissolved in 1X PBS for an additional 36 h at 4 °C, and frozen. Coronal cryostat sections (40 μm) covering the antero-posterior extent of the striatum were used for labeling. Brain sections were permeabilized in PBS with 0.4% Triton X-100 (PBST) for 30 min at room temperature. Sections were then incubated in a blocking solution composed of PBST with 5% bovine serum albumin for 1h at room temperature. Free-floating sections were incubated overnight at 4 °C in mouse anti-TH (1/1000, Millipore, MAB318) or in rabbit anti-RFP (2/1000, Clinisciences, PM005) and goat anti-ChAT (1/100, Millipore, AB144P). After exposure to primary antibody, two different protocols were used to reveal TH or mCherry/ChAT expression. For TH detection, sections were incubated in a PBS solution containing the biotinylated secondary antibody (Biotinylated goat anti-mouse, #115-065-166, Jackson Immunoresearch, diluted 1/200) for 1 hour at room temperature. Immunoperoxidase detection was then performed by enzymatic reaction with an avidin-peroxidase complex for 1 h (Vectastain ABC HRP Kit Peroxidase Standard, Vector Lab.). After washing in PBS, the complex formed was revealed by incubation in 3,3’-Diaminobenzidine tetrahydrochloride (Sigmafast 3,3’-DAB tablets, D4293, Sigma-Aldrich). Following a final wash in PBS, slices were mounted on superfrost slides and left to dry overnight. The next day, the slides were dehydrated and incubated in xylene baths, coverslipped and mounted in synthetic DPX mounting medium (Sigma Aldrich). Quantification of striatal TH immunostaining was performed by digitized image analysis using “Densirag” analysis system (BIOCOM, France). The mean optical density (OD) value was determined from at least two sections per animal distributed on the antero-posterior extent of the dorsal striatum after subtracting the background signal measured in a region lacking dopaminergic terminals (parietal cortex). All the 6-OHDA-injected mice used in this study exhibited > 70% loss of DA axons. To reveal mCherry and ChAT expression, sections were incubated in Alexa Fluor 594 donkey anti-rabbit (1/200, Invitrogen, A31572) and Alexa Fluor 488 donkey anti-goat (1/200, Invitrogen, A11055). Sections were then mounted onto SuperFrost Plus glass slides (VWR) and coverslipped with FluorSave mounting media (Merck Chemicals).

### FACS-sorting

Striatal single cell suspensions were prepared after vehicle or 6-OHDA injected (28 to 31 days post-injection). 250 μm striatal slices were prepared as for *in vitro* electrophysiological recordings. Striata were then microdissected and incubated for 10 minutes in HibernateA minus calcium (Brainbits, HA-Ca) medium supplemented with 25 mM L-Glutamine (Gibco), pre-warmed at 37°C. Striatal pieces were then transferred to 10 ml sterilin tubes containing 3ml of HA-Ca + 25 mM L-Glutamine medium supplemented with 2 mg/mL papaïn (Whortington) and 25 mg/mL dispase (Gibco) and incubated for 30 minutes at 37°C with shaking (200 rpm). After centrifugation for 5 minutes at 1100 rpm (room temperature), digestion medium was discarded and striatal pieces were transferred into 1ml of ice-cold FACS-buffer containing HibernateA plus calcium, no phenol red (Brainbits), 2% (v/v) B27 supplement (Invitrogen) and 0.5 mM L-Glutamine (Gibco). Striatal pieces were mechanically triturated by realizing 7-10 passages through 22G, 23G and 26G needles. In between each type of needle, striatal pieces were allowed to settle for 2 min, 500 μL of suspension was filtered on a 70 μm cell strainer (Gibco), collected in a separate tube and maintained on ice. Then 500 μL of fresh FACS buffer were added to the striatal pieces before resuming the trituration. Finally, cells were precipitated by centrifugation for 20 minutes at 1100 rpm (4°C), medium was discarded and replaced with fresh ice-cold FACS buffer. Sytox Blue (ThermoFisher) viability dye was added to cells suspensions right before FACS. Tdtomato+ MSNs were sorted on a FACS Aria cell sorter (BD Bioscience) by gating on Sytox Blue- (live cells) tdTomato+ cells directly in ice-cold RLT lysis buffer from RNeasy Micro Kit (Quiagen), supplemented with 10% (v/v) β-mercaptoethanol (Sigma), and snap-frozen at −80°C before RNA extraction. During tissue preparation, 2 striatal slices per animal were spared to assess the extent of 6-OHDA lesion by anti-TH immunochemistry.

### RNA extraction

RNA from sorted tdTtomato+ D1 MSNs was extracted using RNeasy Micro Kit (Quiagen), according to manufacturer’s instructions. For quality and quantity determination, 1 μL of extracted RNAs was loaded on a Agilent Pico 6000 chip and ran on the 2100 Bioanlayzer system (Agilent).

### Reverse Transcription followed by quantitative polymerase chain reaction (RT-qPCR)

2,2 ng of high-quality total RNA (RIN > 8.5) were reverse-transcribed into cDNA using the Superscript III enzyme (ThermoFisher) and used as template for qPCR. Specific primers for the different genes of interest were designed using either the Universal ProbeLibrary (ProbeFinder version 2.5 for mouse, Roche Diagnostics) or PrimerBlast (NCBI) algorythm and chosen intron-spanning regions. Quantification of gene expression was done using Syber Green (ROX) master mix (Thermofisher) and the StepOne Real-Time PCR apparatus (ThermoFisher). Amplification of single PCR products was confirmed by the melting curves. The relative quantity of transcript encoding each gene vas determined by normalization to *Gapdh*, using the standard delta Ct method. The following primers were used:

- *Chrma1*-Fw: AAGATGGATTGAATGAGGCTGC, *Chrma1*-Rse: CCTCCAGTCACAAGATTTTTCTCA,
- *Chrma4*-Fw: CAGCGGAGCAAGACAGAAG, *Chrma4*-Rse: CATTGACAGGTGTGAAGTTCG,
- *Gapdh*-Fw: ATGGTGAAGGTCGGTGTGA, *Gapdh*-Rse: AATCTCCACTTTGCCACTGC.

### Rotarod

The apparatus (LE8205, Bioseb) consisted of a rod (30 mm in diameter) suspended horizontally at a height of 20 cm from the floor. Two days before starting the experiments, mice were placed for a few moments on the non-rotating cylinder to familiarize them with the experimental environment. Four trials per day were then performed for 5 days at constant speed (12 rpm) and the latency to fall from the rod was measured with a cut-off time of 60 s. Ten days after this training period, mice were tested for 3 days with accelerated speed, from 4 to 40 rpm over 300 s. Trials were spaced 15 min apart. Rotarod were performed 25 to 28 days after 6-OHDA and/or DREADD injections.

### Quantification and statistical analysis

Data analysis was performed with Clampfit 10.7 (Molecular Devices, Inc.) and Spike 2 (Cambridge Electronic Design Limited, Cambridge, UK). The statistical analyses were two-tailed statistical tests with a risk α set at 0.05 and were performed in GraphPad Prism 9 software (San Diego, CA, USA). Within a set of experimental data, the paired t-test was used for dependent data excepts if the normality distribution test failed (Shapiro-Wilk test, p < 0.05). In the latter case, the Wilcoxon signed rank test was used. For independent data, we applied the normality (Shapiro-Wilk test) and equal variance tests. A t-test was used if the distributions were normal and the group variances were equal. Otherwise, the Mann-Whitney signed rank test was used.

## SUPPLEMENTAL TABLES

**Table S1.**
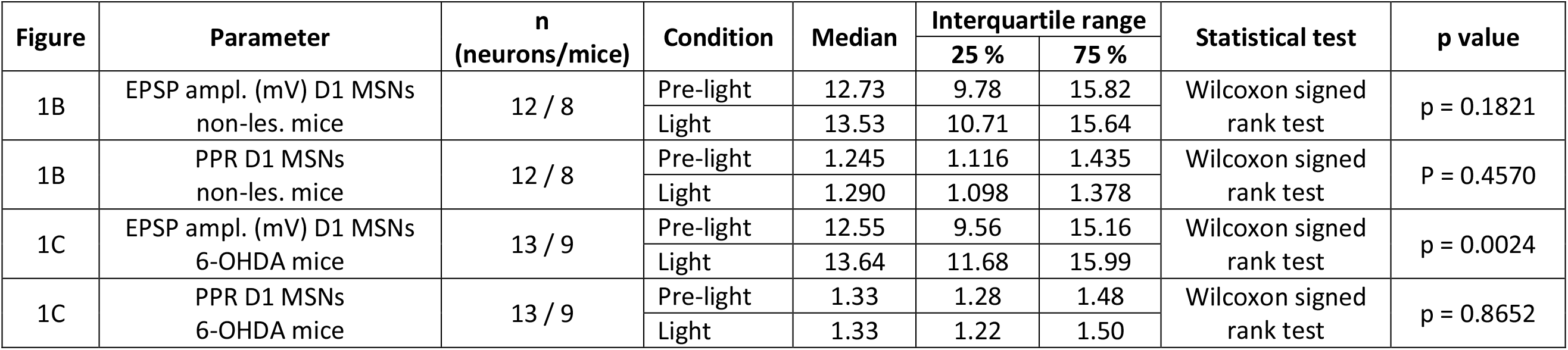
Related to Figure 1.

| Figure | Parameter | n<br>(neurons/mice) | Condition | Median | Interquartile range |  | Statistical test | p value |
| --- | --- | --- | --- | --- | --- | --- | --- | --- |
|  |  |  |  |  | 25 % | 75 % |  |  |
| 1B | EPSP ampl. (mV) D1 MSNs<br>non-les. mice | 12 / 8 | Pre-light | 12.73 | 9.78 | 15.82 | Wilcoxon signed<br>rank test | p = 0.1821 |
|  |  |  | Light | 13.53 | 10.71 | 15.64 |  |  |
| 1B | PPR D1 MSNs<br>non-les. mice | 12 / 8 | Pre-light | 1.245 | 1.116 | 1.435 | Wilcoxon signed<br>rank test | P = 0.4570 |
|  |  |  | Light | 1.290 | 1.098 | 1.378 |  |  |
| 1C | EPSP ampl. (mV) D1 MSNs<br>6-OHDA mice | 13 / 9 | Pre-light | 12.55 | 9.56 | 15.16 | Wilcoxon signed<br>rank test | p = 0.0024 |
|  |  |  | Light | 13.64 | 11.68 | 15.99 |  |  |
| 1C | PPR D1 MSNs<br>6-OHDA mice | 13 / 9 | Pre-light | 1.33 | 1.28 | 1.48 | Wilcoxon signed<br>rank test | p = 0.8652 |
|  |  |  | Light | 1.33 | 1.22 | 1.50 |  |  |

**Table S2.** Related to Figures S2, S3 and S4.

| Figure | Parameter | n<br>(neurons/mice) | Condition | Median | Interquartile range |  | Statistical test | p value |
| --- | --- | --- | --- | --- | --- | --- | --- | --- |
|  |  |  |  |  | 25 % | 75 % |  |  |
| S2A | EPSP ampl. (mV) D1 MSNs<br>non-les. mice + neostigmine | 6 / 5 | Pre-light | 8.12 | 5.93 | 11.70 | Paired t-test | p = 0.9893 |
|  |  |  | Light | 8.23 | 5.14 | 12.57 |  |  |
| S2A | PPR D1 MSNs<br>non-les. mice + neostigmine | 6 / 5 | Pre-light | 1.295 | 1.160 | 1.353 | Paired t-test | p = 0.6838 |
|  |  |  | Light | 1.280 | 1.203 | 1.400 |  |  |
| S3B | EPSP ampl. (mV) MSNs<br>non-les. mice / in vivo | 10 / 6 | Pre-light | 10.47 | 6.05 | 13.63 | Wilcoxon signed<br>rank test | p = 0.5566 |
|  |  |  | Light | 9.77 | 6.30 | 12.95 |  |  |
| S3B | PPR MSNs<br>non-les. mice / in vivo | 10 / 6 | Pre-light | 1.770 | 1.503 | 2.556 | Wilcoxon signed<br>rank test | p = 0.3750 |
|  |  |  | Light | 1.692 | 1.595 | 2.657 |  |  |
| S4 | EPSP ampl. (mV) D1 MSNs<br>6-OHDA mice / no opsin | 18 / 5 | Pre-light | 12.05 | 9.10 | 14.83 | Wilcoxon signed<br>rank test | p = 0.2381 |
|  |  |  | Light | 11.45 | 9.18 | 17.13 |  |  |
| S4 | PPR D1 MSNs<br>6-OHDA mice / no opsin | 18 / 5 | Pre-light | 1.250 | 1.000 | 1.400 | Wilcoxon signed<br>rank test | p = 0.1558 |
|  |  |  | Light | 1.300 | 1.100 | 1.500 |  |  |

**Table S3.** Related to Figure 2 and S6.

| Figure | Parameter | n<br>(neurons/mice) | Condition | Median | Interquartile range |  | Statistical test | p value |
| --- | --- | --- | --- | --- | --- | --- | --- | --- |
|  |  |  |  |  | 25 % | 75 % |  |  |
| 2A | sIPSC frequency (Hz) | 7 / 2 | Non-les.<br>mice | 1.72 | 1.50 | 1.92 | Unpaired t-test | p = 0.1005 |
|  |  | 10 / 5 | 6-OHDA<br>mice | 1.28 | 1.03 | 1.52 |  |  |
| 2B | qPCR: relative expression<br>*Chrma1* (arbitrary unit) | na / 7 | Non-les.<br>mice | 0.02054 | 0.01873 | 0.02256 | Mann-Whitney<br>signed rank test | p = 0.5350 |
|  |  | na / 7 | 6-OHDA<br>mice | 0.02193 | 0.02100 | 0.02240 |  |  |
| 2B | qPCR: relative expression<br>*Chrma4* (arbitrary unit) | na / 7 | Non-les.<br>mice | 0.02335 | 0.02051 | 0.02509 | Mann-Whitney<br>signed rank test | p = 0.1649 |
|  |  | na / 7 | 6-OHDA<br>mice | 0.03044 | 0.02152 | 0.03247 |  |  |
| S6B | qPCR: Ct values *Gapdh* | na / 7 | Non-les.<br>mice | 21.84 | 21.72 | 22.62 | Mann-Whitney<br>signed rank test | p = 0.7104 |
|  |  | na / 7 | 6-OHDA<br>mice | 21.77 | 21.62 | 23.25 |  |  |
| 2F | EPSP ampl. (mV) D1 MSNs<br>6-OHDA mice + tropicamide | 10 / 2 | Pre-light | 13.11 | 10.54 | 14.95 | Wilcoxon signed<br>rank test | p = 0.3223 |
|  |  |  | Light | 13.68 | 10.55 | 15.66 |  |  |
| 2F | PPR D1 MSNs<br>6-OHDA mice + tropicamide | 10 / 2 | Pre-light | 1.385 | 1.235 | 1.620 | Wilcoxon signed<br>rank test | p = 0.0820 |
|  |  |  | Light | 1.355 | 1.205 | 1.563 |  |  |
| 2F | EPSP ampl. (mV) D1 MSNs<br>6-OHDA mice + PKI | 8 / 2 | Pre-light | 13.55 | 8.25 | 14.05 | Wilcoxon signed<br>rank test | p = 0.8672 |
|  |  |  | Light | 13.75 | 7.88 | 16.28 |  |  |
| 2F | PPR D1 MSNs<br>6-OHDA mice + PKI | 8 / 2 | Pre-light | 1.345 | 1.223 | 1.615 | Wilcoxon signed<br>rank test | p = 0.2969 |
|  |  |  | Light | 1.365 | 1.275 | 1.590 |  |  |

**Table S4.** Related to Figure 3.

| Figure | Parameter | n<br>(neurons/mice) | Condition | Mean ± SEM | Statistical test | p value |
| --- | --- | --- | --- | --- | --- | --- |
| 3B | EPSP ampl. (mV)<br>non-les. mice - Average | 6 / 6 | Pre-pairing | 11.03 ± 1.24 | Two-tailed<br>Paired t test | p = 0.0457 |
| | | | Post-pairing (20-30 min) | 13.75 ± 0.72 | | |
| 3B | EPSP ampl. (mV)<br>6-OHDA mice - Average | 5 / 5 | Pre-pairing | 10.51 ± 1.57 | Two-tailed<br>Paired t test | p = 0.5642 |
| | | | Post-pairing (20-30 min) | 10.03 ± 2.02 | | |
| 3B | EPSP ampl. (mV) 6-OHDA<br>mice + CIN inhib. - Average | 5 / 5 | Pre-pairing | 9.48 ± 0.47 | Two-tailed<br>Paired t test | p = 0.6223 |
| | | | Post-pairing (20-30 min) | 10.40 ± 1.37 | | |
| 3C | EPSP ampl. (mV)<br>non-les. mice<br>- Individual cells | 6 / 6 | Cell 1 pre-pairing | 7.60 ± 0.31 | Two-tailed<br>Paired t-test | p = 0.0014 |
| | | | Cell 1 post-pairing | 11.09 ± 0.96 | | |
| | | | Cell 2 pre-pairing | 7.96 ± 0.78 | Two-tailed<br>Paired t-test | p < 0.0001 |
| | | | Cell 2 post-pairing | 14.26 ± 0.51 | | |
| | | | Cell 3 pre-pairing | 12.51 ± 0.74 | Two-tailed<br>Paired t-test | p = 0.0084 |
| | | | Cell 3 post-pairing | 16.03 ± 0.92 | | |
| | | | Cell 4 pre-pairing | 9.76 ± 0.31 | Two-tailed<br>Paired t-test | p = 0.007 |
| | | | Cell 4 post-pairing | 11.34 ± 0.45 | | |
| | | | Cell 5 pre-pairing | 14.91 ± 0.40 | Two-tailed<br>Paired t-test | p = 0.5712 |
| | | | Cell 5 post-pairing | 14.54 ± 0.50 | | |
| | | | Cell 6 pre-pairing | 13.14 ± 0.76 | Two-tailed<br>Paired t-test | p = 0.2243 |
| | | | Cell 6 post-pairing | 14.48 ± 0.78 | | |
| 3C | EPSP ampl. (mV)<br>6-OHDA mice<br>- Individual cells | 5 / 5 | Cell 1 pre-pairing | 12.37 ± 0.59 | Two-tailed<br>Paired t-test | P = 0.0004 |
| | | | Cell 1 post-pairing | 9.17 ± 0.37 | | |
| | | | Cell 2 pre-pairing | 15.51 ± 0.88 | Two-tailed<br>Paired t-test | p = 0.1883 |
| | | | Cell 2 post-pairing | 17.06 ± 0.76 | | |
| | | | Cell 3 pre-pairing | 7.40 ± 0.59 | Two-tailed<br>Paired t-test | p = 0.0336 |
| | | | Cell 3 post-pairing | 5.95 ± 0.30 | | |
| | | | Cell 4 pre-pairing | 6.44 ± 0.62 | Two-tailed<br>Paired t-test | p = 0.0839 |
| | | | Cell 4 post-pairing | 7.62 ± 0.26 | | |
| | | | Cell 5 pre-pairing | 8.10 ± 0.63 | Two-tailed<br>Paired t-test | p = 0.124 |
| | | | Cell 5 post-pairing | 9.19 ± 0.29 | | |
| 3C | EPSP ampl. (mV)<br>6-OHDA mice + CIN inhib.<br>- Individual cells | 5 / 5 | Cell 1 pre-pairing | 9.27 ± 0.40 | Two-tailed<br>Paired t-test | p = 0.4446 |
|  |  |  | Cell 1 post-pairing | 9.71 ± 0.55 |  |  |
|  |  |  | Cell 2 pre-pairing | 7.45 ± 0.28 | Two-tailed<br>Paired t-test | p = 0.0002 |
|  |  |  | Cell 2 post-pairing | 10.94 ± 0.95 |  |  |
|  |  |  | Cell 3 pre-pairing | 9.53 ± 0.76 | Two-tailed<br>Paired t-test | p < 0.0001 |
|  |  |  | Cell 3 post-pairing | 14.77 ± 0.89 |  |  |
|  |  |  | Cell 4 pre-pairing | 8.29 ± 0.95 | Two-tailed<br>Paired t-test | p = 0.3167 |
|  |  |  | Cell 4 post-pairing | 7.34 ± 0.42 |  |  |
|  |  |  | Cell 5 pre-pairing | 8.74 ± 0.25 | Two-tailed<br>Paired t-test | p < 0.0001 |
|  |  |  | Cell 5 post-pairing | 6.46 ± 0.19 |  |  |

**Table S5.**
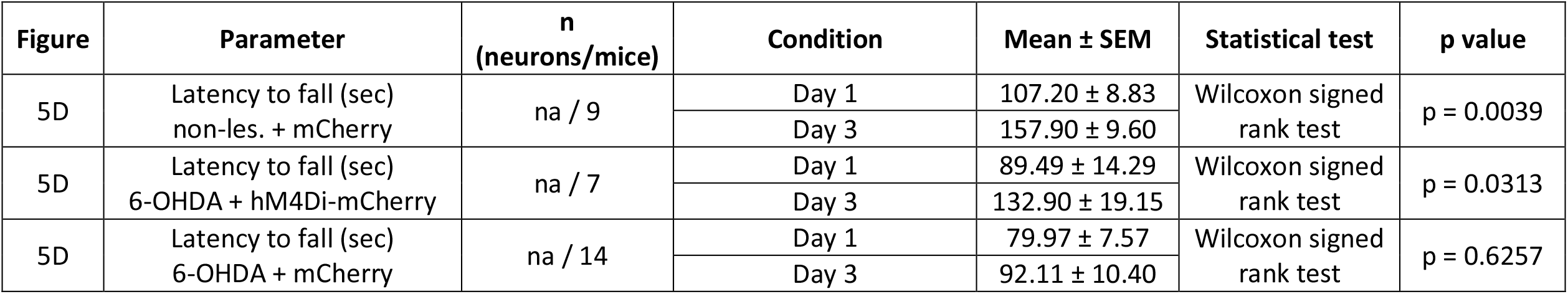
Related to Figure 4.

| Figure | Parameter | n<br>(neurons/mice) | Condition | Mean ± SEM | Statistical test | p value |
| --- | --- | --- | --- | --- | --- | --- |
| 5D | Latency to fall (sec)<br>non-les. + mCherry | na / 9 | Day 1 | 107.20 ± 8.83 | Wilcoxon signed<br>rank test | p = 0.0039 |
|  |  |  | Day 3 | 157.90 ± 9.60 |  |  |
| 5D | Latency to fall (sec)<br>6-OHDA + hM4Di-mCherry | na / 7 | Day 1 | 89.49 ± 14.29 | Wilcoxon signed<br>rank test | p = 0.0313 |
|  |  |  | Day 3 | 132.90 ± 19.15 |  |  |
| 5D | Latency to fall (sec)<br>6-OHDA + mCherry | na / 14 | Day 1 | 79.97 ± 7.57 | Wilcoxon signed<br>rank test | p = 0.6257 |
|  |  |  | Day 3 | 92.11 ± 10.40 |  |  |

## Notes

### Competing Interest Statement

The authors have declared no competing interest.

